# A calcium imaging pipeline to detect and quantify compound-specific effects in human and mouse astrocytes and astrocyte-neuron cocultures

**DOI:** 10.64898/2026.03.19.712916

**Authors:** Jeremy Krohn, Larissa Breuer, Susanne Wegmann, Camin Dean

## Abstract

Astrocytes are crucial mediators of diverse aspects of brain function, including energy metabolism and synapse formation and maturation. Calcium is the primary information carrier in astrocytes, reporting cellular health and activity, and can be measured using fluorescent indicators. However, this readout is not yet widely used to screen and evaluate disease models and drug candidates. Here, we adapted a simple automated calcium imaging pipeline with key output parameters that characterize changes in astrocytic calcium signaling. We compared calcium responses in mouse astrocyte monocultures and astrocyte-neuron cocultures using GFAP-driven membrane-targeted GCaMP6f, with human astrocytes differentiated from two different induced pluripotent stem-cell lines using the calcium dye Cal520-AM. Event-based analysis reported similarities and differences in mean fluorescence, amplitude, frequency, duration, and area of calcium responses. We benchmarked the pipeline using the purinergic receptor agonist ATP to increase astrocyte activity, and the ER calcium pump blocker CPA to decrease activity across all culture models. Glutamatergic and serotonergic receptor function was tested with glutamate and lysergic acid diethylamide (LSD). LSD decreased activity in mouse cocultured astrocytes, but increased activity in human astrocytes. Furthermore, the addition of human recombinant Tau oligomers, an in vitro model of Alzheimer’s disease pathology, decreased activity in both mouse and human astrocytes. This pipeline can be used to quickly and easily characterize effects of astrocyte-targeting compounds, effects of non-astrocyte-targeting compounds on astrocyte activity, and rescue of disease models that affect astrocyte function, in mouse and human astrocytes and astrocyte-neuron cocultures.

## INTRODUCTION

Astrocytes are the second largest cell population in the brain. They contribute to the formation of the blood-brain barrier (Alvarez et al., 2013), supply neurons with metabolites (Bolaños & Magistretti, 2025), and facilitate synapse growth, maturation, and plasticity (Chung et al., 2015; Fossati et al., 2020). Despite their critical roles, investigation of astrocyte function remains underrepresented in disease research and drug development. Drug discovery pipelines often follow a neurocentric approach. Consequently, the effect of pharmacological agents on astrocytes (either positive or negative) is often unknown at early stages. For example, Fluoxetine, a serotonin reuptake inhibitor used to treat depressive disorder, elicits calcium responses in astrocytes independent of neuronal activity (Schipke et al., 2011) and can inhibit the switch to reactive A1 astrocytes in a depression model (Fang et al., 2022). Memantine, an NMDAR antagonist used in Alzheimer’s disease, was shown to cause the release of neurotrophic factors from astrocytes (Wu et al., 2009), which may be beneficial. However, Haloperidol, a dopamine receptor antagonist used to treat schizophrenia, causes calcium-dependent cytotoxicity in human astrocytes (Hsu & Liang, 2020), actively harming patients. Testing the effect of compounds on astrocytes is therefore essential for the development of novel, and improvement of existing, therapeutic approaches targeting both astrocytic and neuronal function.

Calcium is a key messenger in astrocytes, associated with astrocyte function and astrocytic modulation of neuronal activity. Calcium signaling can be used as an indicator of cellular activity or dysfunction in diseases. For example, a mouse model of Huntington’s disease displays reduced amplitude and frequency of spontaneous calcium events (Jiang et al., 2016), and mouse astrocyte cultures treated with oligomeric recombinant human Tau have less frequent ATP-induced calcium transients with reduced mean fluorescence (Piacentini et al., 2017). Furthermore, treatment of cultures with β-amyloid peptides increases calcium activity in astrocytes, but not neurons (Abramov et al., 2003).

Astrocyte monocultures can be used to directly test astrocyte-specific mechanisms to treat neurological diseases and disorders, as well as to test toxicity and off-target effects of compounds designed to target neurons. However, astrocyte monocultures have less physiological morphology and function in the absence of neurons. In coculture with neurons, astrocytes become ramified (Martinez-Lozada et al., 2023) and, together with neurons, form tripartite synapses (Araque et al., 1999). Astrocytes sense transmitters such as glutamate (Cai et al., 2000; James et al., 2011; Porter & McCarthy, 1996) or serotonin (Porter & McCarthy, 1997; Whitaker-Azmitia et al., 1993) released during neuronal activity and respond with calcium elevations and the release of neuroactive transmitters to modulate synaptic activity (Perea & Araque, 2005; Porter & McCarthy, 1996). Using both monocultures and cocultures could improve drug discovery by identifying direct vs. indirect effects of candidate compounds on astrocytes, while also testing for beneficial or detrimental effects.

The translation of insights from murine models to human patients often fails (Atkins et al., 2020; Bailey et al., 2014; Pound & Ritskes-Hoitinga, 2018; van Meer et al., 2012; Van Norman, 2019). In the case of astrocytic targets (or testing for side effects in astrocytes), it is especially important to test human systems given the vast physiological differences between rodent and human astrocytes. For example, astrocytes are less conserved across species than other cell types in terms of gene expression (Hodge et al., 2019; Zhang et al., 2016) and disease risk genes (Penney et al., 2020). Functionally, calcium transients in primary human astrocytes are faster than those observed in mice (Oberheim et al., 2009; Zhang et al., 2016). Human-derived model systems have been shown to better predict successful clinical trials (Voskoglou-Nomikos et al., 2003). Using ELISA, MTT, and apoptosis assays, human astrocytes have been used to screen for inhibitors of toxic astrogliosis (Masvekar et al., 2022), compounds protecting against oxidative stress (Thorne et al., 2016), and neurotoxic compounds (Pei et al., 2016). We propose that astrocytic calcium signals can provide a functional readout of drug activity and cell health, especially if used in a standardised, validated pipeline.

Most studies on astrocyte calcium imaging focus on a specific output parameter, such as amplitude (Allen et al., 2022) or event duration (Anding et al., 2025), in a single model system, but a comprehensive overview of different parameters is often missing. Here we sought to adapt and refine a simple calcium imaging analysis pipeline that reliably captures any and all potential changes in astrocytic calcium signaling. We integrated semi-automated microscopy-based astrocyte calcium imaging with AQuA event-based detection software (Wang et al., 2019) into an automated analysis pipeline that outputs multiple fundamental parameters of astrocytic calcium activity to fully characterize astrocytic effects in diverse model systems. We compared calcium dynamics in mouse astrocytes in the absence and presence of neurons, and in human astrocytes derived from two induced pluripotent stem cell (iPSC) lines and identified compound-specific effects on a set of core parameters: mean fluorescence, amplitude, frequency, duration, and area of calcium events. We benchmarked the analysis using ATP and CPA, and tested compounds targeting glutamate receptors (glutamate and MPEP) and serotonergic receptors (LSD). Comparison of calcium dynamics in mouse and human astrocytes revealed striking differences in response to LSD treatment. In addition, treatment with recombinant human Tau oligomers served as an example disease model, with simultaneous recording of neuronal and astrocytic calcium signals. Together, the parameters reported by our pipeline provide a comprehensive overview of calcium signaling dynamics in mouse and human astrocytes and can therefore be used to determine astrocytic effects of compounds, disease models, and their rescue, to facilitate drug discovery.

## METHODS

### Ethics Statements

This study was conducted in accordance with the principles of the German Animal Welfare Act (Tierschutzgesetz) and guidelines of the GV-SOLAS and FELASA. All procedures involving mice were approved by the local animal welfare officer under the approval number T-CH 0013/20.

Mice of the strain C57B6/J used in this research were housed and cared for in accordance with the standards established by the European Union Directive 2010/63/EU on the protection of animals used for scientific purposes. Mice were housed in groups of 4-5 in standard IVC racks, and had ad libitum access to food and water in a 12/12-hour light-dark cycle at 22±2°C.

The human iPSC-derived astrocytes used in this work were generated in accordance with ethical standards and regulations governing human cell research by bit.bio (ioAstrocytes) or by the BIH Core Unit pluripotent Stem Cells and Organoid (iAstrocytes). The use of these cells complies with the principles outlined in the Declaration of Helsinki and applicable local laws regarding the use of human biological materials.

### Mouse astrocytic culture

Cultures enriched for astrocytes were prepared from postnatal day 1-5 mouse pups. The skin was carefully disinfected with ethanol before decapitation. Brains were extracted in ice-cold dissection medium (HBSS (Gibco, 14170-088) containing 10 mM HEPES (Gibco, 15630-056)), the meninges were removed, and cortices were washed three times with dissection medium before dissociation in 0.25% Trypsin-EDTA (Gibco, 25200-056) for 20 minutes at 37°C. Tissue was again washed with dissection medium and then resuspended in prewarmed (37°C) astrocyte culture medium (cDMEM; DMEM, 10% fetal bovine serum (Sigma, F0804 or Gibco, A5256801), 100 U/mL penicillin/streptomycin (Gibco, 15140-122)). After trituration using a 1000 µL pipette tip, the cell number was counted and calculated using a hemacytometer. Dead cells were excluded using a Trypan Blue (Sigma, T8154) counterstain. Cells were plated in T75 flasks (Sarstedt, 83.3911.002) coated with a collagen-polyornithine mix containing 0.12 µg/mL polyornithine (Sigma, P8638), 0.66% (v/v) Collagen type I (Sigma, C3867), 0.06% (v/v) acetic acid (Roth, 3738.4) (protocol adapted from (Schiweck et al., 2021)) at a density of 13,000-26,000 cells/cm². Medium was exchanged the following day and then once per week. Cells were split 1:2 if confluency exceeded 80%. Astrocyte cultures were used until an age of eight weeks and/or three passages. For experiments, astrocytes were split and plated on 96-well glass-bottom plates (Cellvis, P96-1.5H-N) coated with 0.5 mg/mL poly-D-lysine (Sigma, P7886) in 0.1 M borate buffer (Sigma, B9645) at ∼47,000 cells/cm² in cDMEM. Cells were cultured in 200 µL culture medium at 37°C and 5% CO_2_. Half of the medium was exchanged once a week. Cells were used for experiments between 10 and 15 days in vitro.

### Mouse hippocampal culture of astrocytes and neurons

Timed pregnant mice were sacrificed at E18 using CO_2_ and cervical dislocation. The mouse was placed on its back, the abdomen disinfected with ethanol, and skin and muscles dissected to expose the uterus. Embryos were extracted from amniotic sacs and decapitated. Heads were transferred to ice-cold dissection medium, fixed on the rostral side, and a small incision was made along the midline through the skin and skull to extract the brains. Brains were transferred to ice-cold dissection medium, meninges removed, and hemispheres separated. Using bent forceps, hippocampi were dissected, washed three times with dissection medium, and incubated in 0.05% Trypsin-EDTA (Gibco, 25300-054) for 20 minutes at 37°C. Tissue was washed three times in dissection medium and resuspended in cDMEM before it was triturated using a 1000 µL pipette tip. Cell number was counted using a hemacytometer, and dead cells excluded using a Trypan Blue counterstain. Cells were plated on poly-D-lysine-coated glass bottom well plates (24-well (Cellvis, P24-1.5H-N): ∼52,000 cells/cm², 500 µL culture medium; 96-well: ∼47,000 cells/cm², 200 µL culture medium; 384-well (Cellvis, P384-1.5H-N): ∼120,000 cells/cm², 100 µL culture medium) in cDMEM. Medium was changed to cNB containing Neurobasal (Gibco, 21103-049), 1% GlutaMax (Gibco, 35050-038), 100 U/mL penicillin/streptomycin, and 2% B27 (Gibco, 17504-044) the following day. Cells were cultured at 37°C and 5% CO_2_. Half of the medium was exchanged once a week. Cells were used for experiments between 14 and 21 days in vitro.

### Human astrocyte culture (iAstrocytes)

The human iPSC-line BIHi005-A-1E (Breuer et al., 2026) was generated by the Berlin Institute of Health Core Unit pluripotent Stem Cells and Organoids. Pre-differentiated stem cells were induced to an astrocytic fate using doxycycline-controlled expression of NFiB and Sox9. Culture vessels were sequentially coated with 5 µg/mL vitronectin (Sigma, A14700) for 1 hour at 37°C and then 2 µg/mL Biolaminin 521 (Biolamina, LN521) for 2 hours at 37°C, both diluted in calcium and magnesium-free DPBS (Gibco, 14190-144). Coating was removed right before plating without any further washes. Cells were thawed, resuspended, and plated in FGF-medium containing Neurobasal, 1% non-essential amino acids (Gibco, 11140-035), 1% GlutaMax, 1% fetal bovine serum, 2% B27, 8 ng/mL FGF (Peprotech, 100-18B), 5 ng/mL CNTF (Peprotech, 450-13), 10 ng/mL BMP-4 (Peprotech, 120-05ET), including 3 µg/mL doxycycline (Sigma, D3447) and 10 µM ROCK inhibitor (SelleckChem, 51049). Cells were plated on 96-well glass bottom plates at 47,000 cells/cm² and cultured in 200 µL culture medium at 37°C and 5% CO_2_.

After 24 hours, on DIV1, the medium was replaced with FGF-medium without ROCK inhibitor. On DIV2, medium was changed to maturation medium containing 1:1 DMEM-F12 (Gibco, 11330-032) and Neurobasal, 1% GlutaMax, 1% sodium pyruvate (Gibco, 11366-039), 1% N2 supplement (Gibco, 17502-048), 5 ng/mL N-Acetyl-Cysteine (Sigma, A8199), 5 ng/mL heparin-binding EGF-like growth factor (Sigma, E4643), 10 ng/mL CNTF, 10 ng/mL BMP-4, 500 µg/mL dbcAMP (Sigma, D0627), including doxycycline. Half the medium was exchanged every 2-3 days, and cells were used after 21 days in vitro.

### Human astrocyte culture (ioAstrocytes)

Human ioAstrocytes were pre-differentiated and acquired from bit.bio (bit.bio, ioEA1093 Early Access). Culture vessels were sequentially coated with 5 µg/mL vitronectin for 1 hour at 37°C and 2 µg/mL Biolaminin 521 for 2 hours at 37°C, both diluted in calcium and magnesium-free DPBS. Cells were thawed, resuspended, and plated in ioStab: Neurobasal, 1% non-essential amino acids, 1% GlutaMax, 1% fetal bovine serum, 0.1% 2- mercaptoethanol (Gibco, 21985-023), 2% B27, 8 ng/mL FGF, 5 ng/mL CNTF, 10 ng/mL BMP-4. To facilitate cell survival, the ioStab plating medium was supplemented with 10 µM ROCK inhibitor, and astrocyte differentiation was induced with 1 µg/mL doxycycline. Plating density was ∼47,000 cells/cm² on 96-well glass-bottom plates. Cells were cultured in 200 µL culture medium at 37°C and 5% CO_2_.

Medium was exchanged with ioStab without ROCK inhibitor the next day, on DIV1. On DIV2, medium was exchanged to ioMaintenance medium consisting of 1:1 DMEM-F12 and Neurobasal, 1% GlutaMax, 0.1% 2-mercaptoethanol, 1% sodium pyruvate, 1% N2 supplement, 5 ng/mL N-Acetyl-Cysteine, 5 ng/mL heparin-binding EGF-like growth factor, 10 ng/mL CNTF, 10 ng/mL BMP-4, and 500 µg/mL dbcAMP, including 1 µg/mL doxycycline. Half medium changes were done every 2-3 days, and cells were used for experiments after 14 days in vitro.

### Calcium imaging

Mouse astrocytes in mono- or cocultures were transduced with AAV2/5-GFAP:GCaMP6f-LCK (Addgene 52924, from Baljit Khakh; 2.5x10^13^ vg/mL, used at 0.3 µL (24-well plate), 0.07-0.1 µL (96-well plate), or 0.03 µL (384-well plate) per well) or AAV2/5-GFAP:jRGECO-Lck (plasmid: Addgene 100854, from Douglas Kim & GENIE Project; the synapsin promoter was replaced with a GFAP promoter and virus prepared by the Charité Viral Core Facility; 8.8x10^12^ vg/mL, used at 0.1 µL per well in a 96-well plate). Neurons in mouse cultures were transduced with AAV2/1-hSyn:jRCaMP1a (Addgene 100848, from Douglas Kim & GENIE Project; 1.9x10^13^, used at 0.005-0.008 µL per well in a 96-well plate) or AAV2/1-hSyn:GCaMP6f (Addgene 100837, from Douglas Kim & GENIE Project; 2.0x10^13^, used at 0.002-0.005 µL per well in a 96-well plate). Viruses were added to the cultures at least seven days prior to imaging and remained in the culture with continued half medium changes every 2-3 days.

For human astrocyte calcium imaging, the calcium dye Cal520-AM (AAT Bioquest, 21130) was reconstituted in DMSO (Roth, A994.1) and was added to the culture medium at 1-2 µM for 15-30 minutes at 37°C and 5% CO_2_. Before imaging, the culture medium was replaced with imaging medium. The imaging medium for most experiments was FluoroBrite™ DMEM (Gibco, A1896701), but some Tau and LSD experiments were imaged in BrainPhys™ Imaging Optimized Medium (Stemcell Technologies, 5796).

### Microscopy

Live cell imaging was performed at the Charité facility for Advanced Biomedical Imaging (AMBIO) on a Nikon Widefield Ti2 (Nikon) microscope. The system was equipped with LED illumination (Lumencore, SpectraX) for excitation at 475/28 nm for GFP and 555/28 nm for RGECO. Single-bandwidth emission filters were used to filter the detected signal for the respective wavelengths (GFP: 519/26; mCherry: 595/31). Samples were imaged with a 40x objective (Apo Fluo, air, 0.95 NA) (Nikon), and 2x2 binning was applied to reduce file size and enhance the fluorescent signal-to-noise ratio. Signals were collected with an sCMOS PCO.edge camera (Excelitas) with 300 ms frame intervals for recordings of astrocytes, and 500 ms frame intervals for dual color imaging of astrocytes and neurons. Live-cell imaging experiments were performed at 37°C and 5% CO_2_. Images were acquired with NIS-Elements AR software (Nikon).

### Pharmacology

Baseline imaging was performed in 300 µL (24-well plate), 150 µL (96-well plate), or 60 µL (384-well plate) imaging solution. Compounds were diluted in imaging solution and added to the wells to reach final volumes of 500 µL, 200 µL, or 100 µL, respectively, yielding the final concentrations indicated in Table 1.

**Table 1:** Source and final concentration of compounds tested in astrocyte calcium imaging.

| Compound | Manufacturer | Catalog no. | Dissolved in | Final concentration |
| --- | --- | --- | --- | --- |
| ATP | Tocris | 3245 | ddH<sub>2</sub>O | 50 µM |
| CPA | Sigma | C1530 | DMSO | 20 µM |
| Glutamate | Sigma | 49621 | ddH<sub>2</sub>O | 100 µM |
| MPEP | Tocris | 1212 | DMSO | 50 µM |
| Gabazine | Tocris | 1262 | ddH<sub>2</sub>O | 10 µM |
| LSD | LGC | MM1346.00 | ddH<sub>2</sub>O | 10 µM |
| Human full-length Tau (2N4R) | Susanne Wegmann (prepared as above) |  | ddH<sub>2</sub>O | 25 nM |

Human Tau oligomers were prepared by mixing 50-60 µM recombinant Tau monomers (2N4R isoform) purified from E. coli (Barghorn et al., 2005) with 0.7 mg/ml heparin (Applichem; A3004) in DPBS with 2 mM dithiothreitol and incubating for six days at 37°C. Fresh 2 mM dithiothreitol was added every second day. Formed aggregates were pelleted by ultracentrifugation (30 minutes at 80,000 x G), resuspended in DPBS and broken into oligomers by sonication, then aliquoted, flash frozen, and stored at -80°C until use.

### Astrocyte signal detection with AQuA

Astrocyte signals were detected using the Astrocyte Quantitative Analysis (AQuA) software package (Wang et al., 2019) (version from September 2023) in MATLAB (Mathworks Inc., Natick, Massachusetts; Version: 9.14.0.2337262 (R2023a) Update 5).

Raw microscopy data were first converted to TIF files using Python (Python Software Foundation, https://www.python.org/, version 3.10.6). To exclude slight photobleaching and disturbance from stage movement, the first 20-40 frames were removed from recordings. Parameters for event detection were adjusted for each dataset by testing 3-5 recordings for proper signal detection. In rare cases, individual files were analyzed with different parameters when an examination of the detected signal indicated over- or underestimation of events (this was the case between different preparations of mouse cocultured astrocytes for Tau experiments, and for one file in iAstrocyte ATP experiments). MATLAB files were then imported into Python for further analysis and visualization of the data.

### Neuronal signal detection with Python

Neuronal calcium signals were extracted from time-lapse images using a custom script in Python. Since we observed mainly synchronous activity in neurons (Movie 4), we calculated the mean pixel intensity of the full field of view per frame, resulting in a single fluorescence trace per recording. The trace was baseline corrected using the ZhangFit algorithm (BaselineRemoval package) to reduce the effects of photobleaching and background noise. Potential neuronal events were detected using a percentile-based amplitude threshold (typically the 80th–90th percentile of the corrected trace) in combination with a minimum peak prominence to exclude small fluctuations and improve detection. Peaks were identified with Python’s find_peaks function in the SciPy module, and for each detected event, the frame index was recorded for downstream frequency analysis.

### Data analysis

Data analysis of calcium signals was performed in Python. Scripts were developed using ChatGPT (OpenAI) and manually inspected and revised to ensure accuracy.

Acute transient effects were only seen in ATP-treated samples after visual inspection of raw data. We therefore analyzed only the first 30 seconds in ATP samples, and full recordings in all other compound-treated samples.

To calculate mean dF/F, we extracted the dF/F traces for each event from the AQuA detection. The sample mean was calculated by averaging the dF/F values per frame across the recording. Group means were calculated by averaging sample means.

For statistical comparison of frequencies, mean fluorescence intensities, and event duration and area sub-categories, we used the Wilcoxon signed-rank test for paired data, and the Mann-Whitney-U test (two groups) or Kruskal-Wallis test with Bonferroni family-wise error rate (FWER) to control for multiple comparisons (three groups), for independent data.

To assess event area, duration, and amplitude in datasets with paired data, a linear mixed model was used to account for the nested structure (multiple events per sample). The experimental condition was set as a fixed effect. The sample identifier was included as a random effect with both random intercept and slope, allowing each sample to have its own baseline reference and its own condition-specific response, thereby accounting for between-sample variability. Data were log-transformed to improve model fit. Model fitting was performed using restricted maximum likelihood (REML), which provides a less biased random effect variance independent of fixed effects. Model fit was evaluated using residual and quantile-quantile plots.

For datasets with independent groups (comparison of mouse and human astrocytes), data were log-transformed, and the experimental condition was set as a fixed effect. Two groups were modelled using a linear model and fitted using ordinary least squares (OLS). Three-group datasets were analyzed using a linear mixed model. The sample identifier was included as a random effect with random intercept and fitted using restricted maximum likelihood (REML). P-values for pairs in the three-group comparison were adjusted using the Bonferroni FWER to control for multiple comparisons. Model fit was evaluated using residual and quantile-quantile plots.

Model coefficients for area, duration, and amplitude were back-transformed to obtain treatment/control ratios, which were expressed as percent change relative to control. Corresponding 95% confidence intervals (CI) were obtained by back-transforming the model-based CI of the fixed effect. These LMM-derived effect sizes and CI are displayed in black in the forest plots on a logarithmic scale. For all other parameters (event frequency, mean dF/F, and area and duration sub-categories), percent change was calculated on a per-sample basis as:

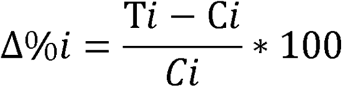

where C*i* and T*i* denote paired control and treatment values from the same sample. The median percent change across samples was used as the central estimate. Uncertainty for these non-LMM metrics was represented using the interquartile range (IQR) of the per-sample percent changes (25^th^–75^th^ percentile). These IQRs reflect between-sample variability in relative treatment effects. In forest plots, both the median percent change and the IQR ranges are displayed in blue.

The experimenter was not blinded to experimental conditions, and no randomization was performed. Statistical significance was assessed at the condition level using an alpha of 0.05.

## RESULTS

### Detection and quantification of astrocytic calcium activity using AQuA

Calcium signals can be used as a functional readout of astrocyte activity. To perform and analyze astrocyte calcium imaging with high throughput, we optimized culture conditions for primary mouse and human iPSC-derived astrocytes cultured in multi-well plates (up to 384 wells), followed by signal detection with AQuA (Wang et al., 2019) and subsequent analysis using custom Python scripts. The detection parameters for AQuA were determined per dataset and kept consistent throughout. Example images demonstrate accurate detection of both spontaneous activity and ATP-induced calcium waves (Fig. 1A, Movie 1). They also serve to verify astrocyte health and morphology, optimal imaging settings for sufficient signal-to-noise, spatial and temporal resolution, detection parameters, and a qualitative overview of event areas. In output images, each colored shape indicates an individual detected event, of which different characteristics, such as the area, duration, peak amplitude, or dF/F can be extracted from AQuA.

**Figure 1:**
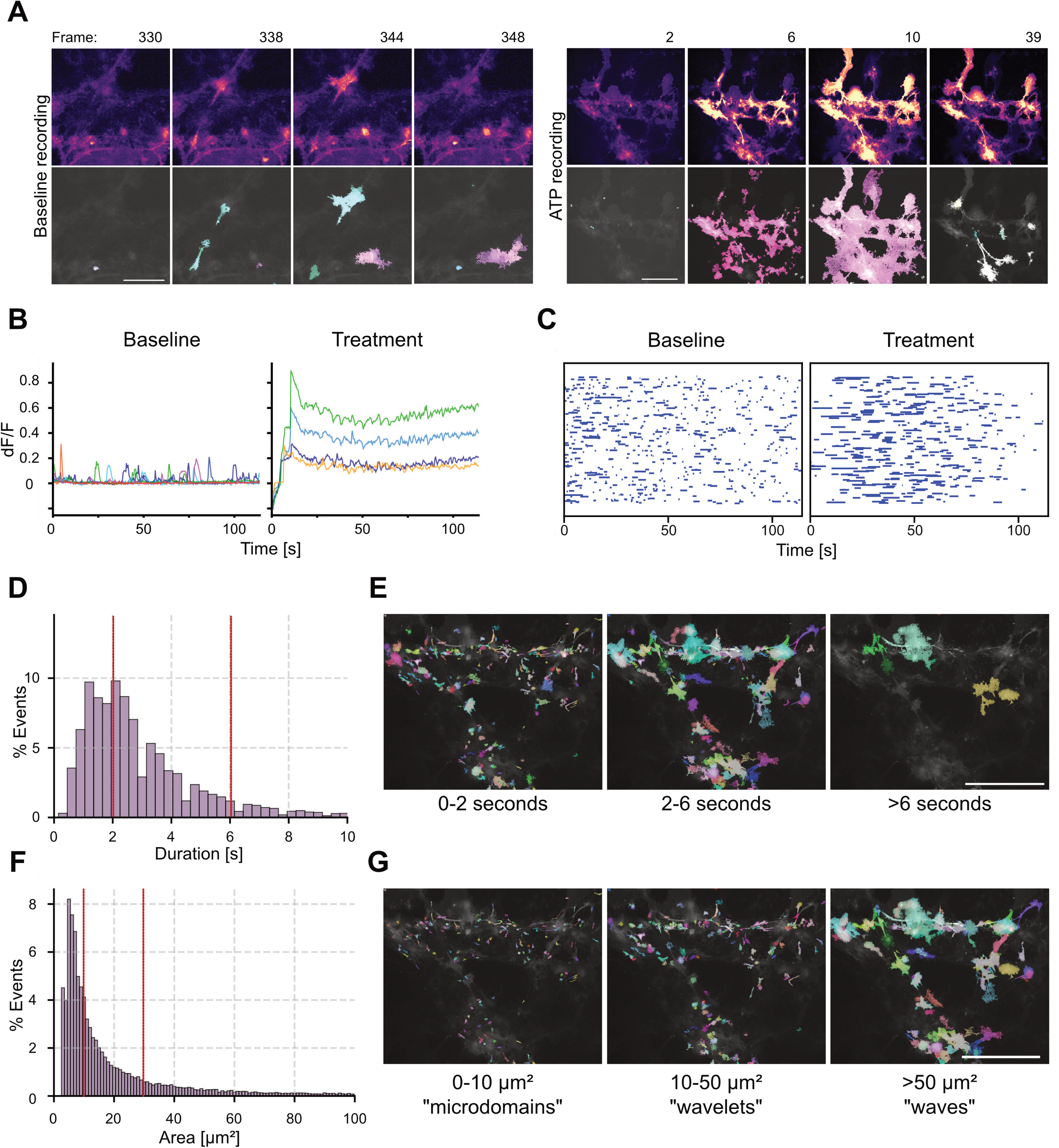
Detection and characterization of astrocytic calcium events. **(A)** Representative images showing raw calcium signals (upper panels) and accurate detection with AQuA (bottom panels) for spontaneous (left) and ATP-induced (right) activity. Frame interval is 300 ms. Scale bars are 100 µm. **(B)** Example event traces from baseline and treatment recordings. Each color corresponds to an individual event and displays fluorescence across the full recording. **(C)** Raster plot showing occurrence of individual astrocytic events as blue lines, with length corresponding to event duration. **(D)** Histogram showing the distribution of event durations in seconds; events longer than 10 seconds (2.6% of events) are not shown. Red lines indicate the subclasses used within this study. **(E)** Representative images with events of the respective duration class overlayed. Scale bar is 100 µm. **(F)** Histogram showing the distribution of event areas in µm²; events larger than 100 µm² (11.5% of events) are not shown. Red lines indicate the subclasses used within this study. **(G)** Representative images with events of the respective size class overlayed. Scale bar is 100 µm.

Fluorescent traces (Fig. 1B) are extracted for each event and display the fluorescence intensity in the event region across the recording. They give a qualitative overview of potential changes in event amplitude, frequency, or duration induced by a compound or perturbation. From the dF/F traces, the mean dF/F is then calculated, where a fluorescence increase or decrease compared to control recordings could reflect a change in amplitude, frequency, or duration of events, or a combination of these parameters. It is important to note that it is also possible for the mean dF/F to remain unchanged, while other parameters are altered; a decrease in frequency and an increase in duration may occur without a change in mean dF/F, for example. The amplitude of events – the maximum single fluorescence value of a given event – is also extracted. The mean dF/F and amplitude may be similar or show differences, depending on the duration of events. Compounds that increase event duration, for example, would increase the mean dF/F, but may cause no change in event amplitude, or even decrease event amplitude. Raster plots (Fig. 1C) display the raw event counts over time per sample, where each blue line corresponds to a single event, the length of which indicates its duration. Changes in the number or length of lines indicate differences in frequency or duration of events. Event frequency (events/min.) and duration are extracted and quantified, in addition to the area of events.

Both duration and area of events encompass large ranges. We therefore subclassified duration and area into three classes derived from morphological features described in the literature, and the distribution of event durations and areas in our dataset. We pooled 159 baseline recordings from 17 different experiments, containing 38,555 individual events, and plotted a histogram for the event duration. We found a minimum duration of 0.3 seconds (corresponding to the frame interval used in imaging) and a maximum of 114.3 seconds, but most events (97.4%) were found in the range of 0-10 seconds (Fig. 1D). Three different classes of event duration were defined. The first class was set to 0-2 seconds (36.7% of detected events) and contained microdomain activity (Di Castro et al., 2011). A second class was set to 2-6 seconds and includes small propagating events (Di Castro et al., 2011), spotty microdomains (Shigetomi et al., 2011), or short-duration somatic, endfeet, and branch events (Shigetomi et al., 2013). In our data, this class included the majority (53.9%) of events. The third class included all events with a duration longer than 6 seconds, made up 9.5% of detected events, and corresponded to somatic events (Wang et al., 2006) or calcium twinkles (Kanemaru et al., 2014). Example images with detected events of each duration class are shown (Fig. 1E).

We then plotted the histogram for the event area (Fig. 1F), which showed a clear right-skewed distribution. The minimum area was 3.03 µm², and the maximum was 5,578.57 µm², but the majority of events (88.5%) were in the range of 3.03-100 µm². We divided event area into three classes of 0-10, 10-50, and >50 µm². In our dataset, 40.6% of events fall into the 0-10 µm² class, which could correspond to small perisynaptic events, for example. Events in this range have been called “spotty microdomain” events (Shigetomi et al., 2013), focal events (Di Castro et al., 2011), or microdomain activity (Srinivasan et al., 2015). We therefore use the term “microdomains” for this type of activity. The second class, 10-50 µm², made up 40.7% of events and would include events in astrocytic processes (Srinivasan et al., 2015) or expanded events (Di Castro et al., 2011). We refer to these events as “wavelets”. The last class includes all events larger than 50 µm², which contain somatic events (Srinivasan et al., 2015) and intercellular waves. In our dataset, the remaining 18.6% of events were larger than 50 µm². Events detected in the respective classes from one sample are shown (Fig. 1G).

### Benchmarking the pipeline using ATP and CPA in mouse astrocytes

We first benchmarked our analysis pipeline with agents known to reliably increase or decrease intracellular calcium in mouse astrocytes. To test increases in astrocytic calcium signals, we used ATP, a universal energy metabolite that also acts as a signaling molecule. ATP activates G-protein coupled receptors such as P2X, P2U, and P2Y on astrocytic membranes (Chen & Chen, 1996; King et al., 1996; Walz et al., 1994) and induces calcium influx in astrocytes (Neary et al., 1988). ATP is also required to generate propagating calcium waves within and between astrocytes (Guthrie et al., 1999) and reports function of astrocytes (Cai et al., 2000; Guthrie et al., 1999; James et al., 2011; Pivneva et al., 2008; Walz et al., 1994; Zur Nieden & Deitmer, 2006).

Astrocyte cultures were transduced with AAV5 GFAP-driven membrane-targeted GCaMP6f. ATP initiated a rapid elevation of calcium, seen as an increase in fluorescence spanning large proportions of the cells (Fig. 2A, Movie 2). Traces of individual event fluorescence (Fig. 2B) showed a rapid, but transient increase. We therefore analyzed the first 30 seconds after ATP application and found an increase in mean fluorescence (*p*<0.001) in response to ATP (Fig. 2C). The sharp increase in fluorescence closely resembles the response to ATP seen in other publications (Pivneva et al., 2008; Zur Nieden & Deitmer, 2006) and thus confirms the expected effect in our cultures. The amplitude of individual events was also increased by ATP (Fig. 2D; β=1.21, 95%-CI: 1.04, 1.39, *p*=0.011). Raster plots (Fig. 2E) indicated increased event density immediately after ATP application, caused by an increase in both frequency in events per minute (Fig. 2F; *p*=0.0068) and duration (Fig. 2G; β=1.54, 95%-CI: 1.16, 2.03, *p*=0.0026), due to a decrease in fast 0-2 second activity (*p*=0.099). Event area was unchanged by ATP (Fig. 2H), likely because ATP initiated a rapid, single event spanning large proportions of cells immediately after application, but this effect did not occur repeatedly. Statistics are summarized in Figure 2I and Table 2.

**Figure 2:**
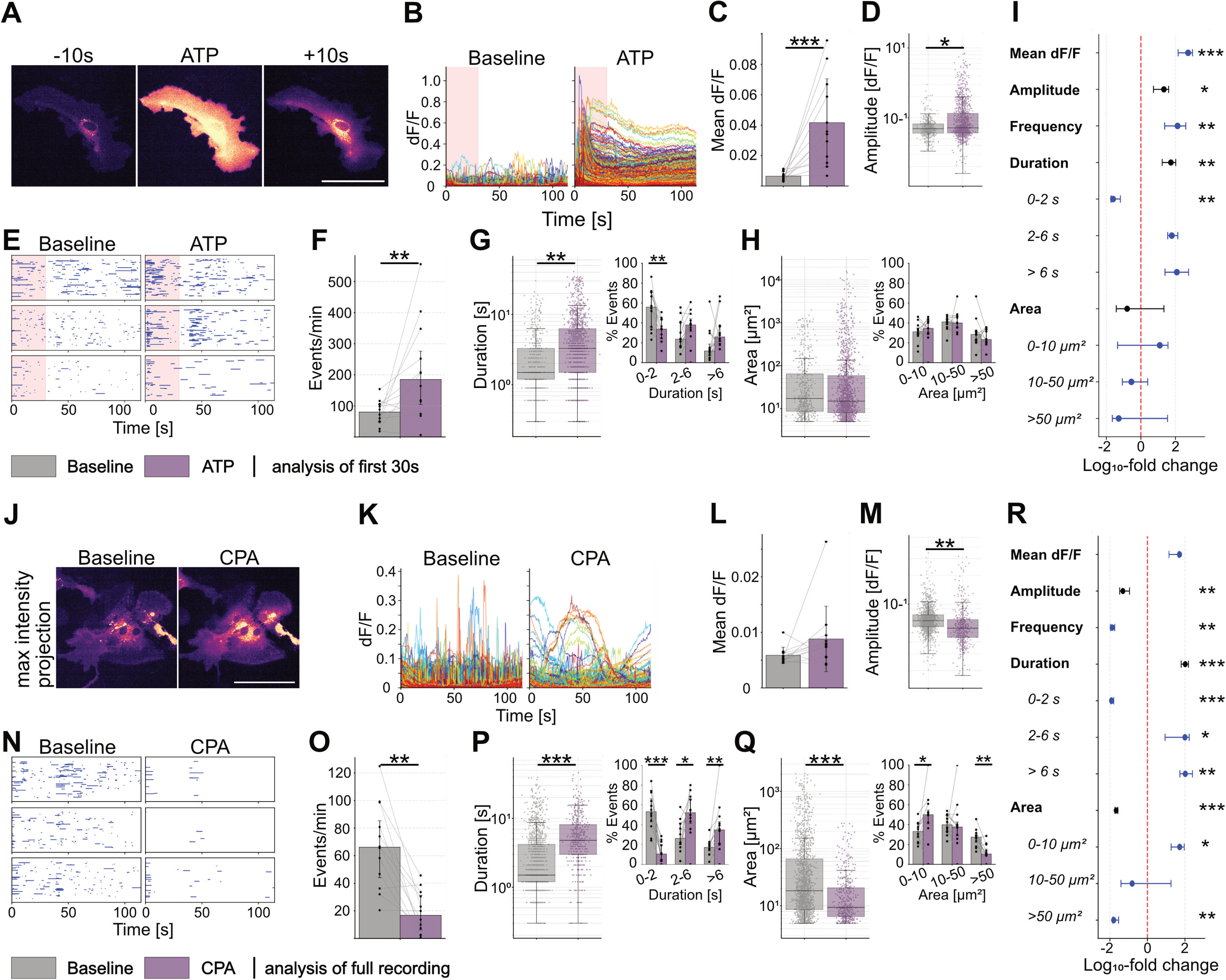
ATP increases, and CPA decreases calcium activity in mouse astrocytes in monocultures. **(A)** Time-lapse images of a typical response induced by 50 µM ATP in mouse astrocyte monocultures transduced with AAV5 GFAP promoter-driven membrane-targeted GCaMP6f. **(B)** Individual event dF/F traces from baseline and treatment recordings. Light red shade indicates 30 second time window used for analysis. **(C)** Mean fluorescence per sample from event traces, displayed as paired data (dots) and group means (bars) ± SD. **(D)** Event amplitude; each dot is an individual event. Boxplots indicate median (center line), interquartile ranges (25^th^-75^th^ percentile), and whiskers extending to 1.5×IQR. **(E)** Raster plots showing occurrence of individual astrocytic events as blue horizontal lines, with length corresponding to event duration. **(F)** Event frequency, displayed as paired regions (dots) and group median (bars) ± IQR. **(G)** Event duration; each dot is an individual event. Boxplots indicate median (center line), interquartile ranges (25^th^-75^th^ percentile), and whiskers extending to 1.5×IQR. Inset (right) shows percent of events per duration subclass paired per-sample (dots) and group median (bars) ± IQR. **(H)** Event area; each dot is an individual event. Boxplots indicate median (center line), interquartile ranges (25^th^-75^th^ percentile), and whiskers extending to 1.5×IQR. Inset (right) shows percent of events per area subclass paired per-sample (dots) and group median (bars) ± IQR. **(I)** Forest plot summarizing treatment effects per parameter as percent change on a log_10_-scale. Amplitude, duration, and area show effect size and confidence intervals derived from linear mixed-effect models in black. Mean dF/F, frequency, and duration and area subclasses, show per-sample paired percent change across all samples as median with interquartile ranges (25^th^-75^th^ percentile) in blue (Wilcoxon signed-rank test). **(J)** Representative max intensity projections, **(K)** event dF/F traces, **(L)** mean fluorescence, **(M)** event amplitude, **(N)** raster plots, **(O)** event frequency, **(P)** event duration and sub-classes (inset), **(Q)** event area and sub-classes (inset), **(R)** and forest plot of mouse astrocytes transduced with AAV5 GFAP promoter-driven membrane-targeted GCaMP6f after 20 µM CPA treatment. n = 12 each for ATP and CPA baseline-treatment pairs; * p > 0.05, ** p > 0.01, *** p > 0.001. Scale bar is 100 µm.

**Table 2:** Descriptive statistics for effects of ATP and CPA in mouse astrocytes in monoculture.

|  | Baseline | ATP | % Change | p-value |
| --- | --- | --- | --- | --- |
| Frequency (events/min); median ± IQR; WSR | 80.00 (49.50–104.50) | 185.00 (104.00–276.00) | 131.25% | 0.006836 |
| Mean dF/F; mean ± SD; WSR | 0.01 ± 0.00 | 0.04 ± 0.03 | 539.10% | 0.000977 |
| Amplitude; median ± IQR; LMM | 0.07 (0.06–0.08) | 0.07 (0.06–0.12) | 20.55% | 0.011086 |
| Duration; median ± IQR; LMM | 1.50 (1.20–3.30) | 3.30 (1.50–6.30) | 53.61% | 0.002599 |
| Duration class 0–2 s; median ± IQR; WSR | 55.93 (35.42–70.91) | 33.75 (23.89–43.45) | -39.66% | 0.009925 |
| Duration class 2–6 s; median ± IQR; WSR | 23.67 (16.50–40.42) | 37.98 (31.31–44.06) | 60.47% | 0.266113 |
| Duration class >6 s; median ± IQR; WSR | 11.81 (7.04–18.71) | 25.88 (21.16–36.57) | 119.22% | 0.151367 |
| Area; median ± IQR; LMM | 17.13 (8.62–64.14) | 14.86 (7.97–58.49) | -5.22% | 0.663280 |
| Area class 0–10 μm<sup>2</sup>; median ± IQR; WSR | 31.00 (28.16–34.85) | 34.55 (29.26–41.91) | 11.45% | 0.733398 |
| Area class 10–50 μm<sup>2</sup>; median ± IQR; WSR | 40.96 (38.13–43.56) | 39.92 (35.47–46.84) | -2.54% | 0.569336 |
| Area class >50 μm<sup>2</sup>; median ± IQR; WSR | 28.42 (22.25–31.43) | 23.28 (19.72–33.76) | -18.11% | 0.423828 |
|  | Baseline | CPA | % Change | p-value |
| Frequency (events/min); median ± IQR; WSR | 66.14 (44.09–85.30) | 16.80 (6.17–31.23) | -74.60% | 0.001465 |
| Mean dF/F; mean ± SD; WSR | 0.01 ± 0.00 | 0.01 ± 0.01 | 50.16% | 0.077148 |
| Amplitude; median ± IQR; LMM | 0.07 (0.06–0.08) | 0.06 (0.04–0.07) | -19.34% | 0.001365 |
| Duration; median ± IQR; LMM | 1.50 (1.20–4.20) | 4.80 (3.00–8.10) | 102.60% | 5.72E-10 |
| Duration class 0–2 s; median ± IQR; WSR | 53.20 (42.81–66.15) | 10.53 (4.38–20.17) | -80.21% | 0.000488 |
| Duration class 2–6 s; median ± IQR; WSR | 26.24 (19.85–41.05) | 52.27 (41.36–66.14) | 99.22% | 0.016113 |
| Duration class >6 s; median ± IQR; WSR | 17.17 (13.69–21.00) | 34.90 (20.79–43.14) | 103.26% | 0.006836 |
| Area; median ± IQR; LMM | 18.31 (8.62–66.19) | 9.37 (6.46–20.65) | -45.46% | 1.48E-15 |
| Area class 0–10 μm<sup>2</sup>; median ± IQR; WSR | 33.05 (26.65–40.93) | 50.00 (40.67–54.71) | 51.31% | 0.042480 |
| Area class 10–50 μm<sup>2</sup>; median ± IQR; WSR | 39.85 (33.12–49.50) | 37.65 (28.68–42.67) | -5.52% | 0.518555 |
| Area class >50 μm<sup>2</sup>; median ± IQR; WSR | 27.41 (22.05–29.73) | 10.44 (9.19–15.23) | -61.92% | 0.004883 |

To benchmark decreases in astrocytic activity, we used cyclopiazonic acid (CPA), which inhibits calcium-dependent ATPases on the sarcoplasmic or endoplasmic reticulum and blocks re-entry of calcium into internal stores and thus depletes astrocytes of calcium (Haustein et al., 2014; Parri & Crunelli, 2003; Zur Nieden & Deitmer, 2006). In contrast, microdomain activity, which is often due to extracellular calcium influx, is only reduced and not blocked completely by CPA (Jiang et al., 2016). In monocultures of mouse astrocytes, addition of CPA (Fig. 2J) appeared to prolong fluorescence increases (Fig. 2K). Since this effect persisted, we analyzed full recordings. We observed a decrease in event amplitude (Fig. 2M; β=0.81, 95%-CI: 0.71, 0.92, *p*=0.0014) after addition of CPA. Raster plots (Fig. 2N) indicated strongly decreased activity, which was due to a reduction in event frequency (Fig. 2O; *p*=0.0015), confirming the primary effect of CPA in our model. Event duration was increased (Fig. 2P; β=2.03, 95%-CI: 1.62, 2.53, *p*<0.001), likely due to fewer short 0-2 s events (*p*<0.001), and more longer events (2-6 seconds: *p*=0.016; >6 seconds: *p*=0.0068). Event area was decreased (Fig. 2Q: β=0.55, 95%-CI: 0.47, 0.63, *p*<0.001) because of an increase in smaller microdomain events (*p*=0.0425) and a decrease in larger waves (*p*=0.0049). Statistics are summarized in Figure 2R and Table 2.

### Astrocyte monocultures and cocultures with neurons differ in baseline and evoked activity

Monocultures allow tests of astrocyte-specific effects of compounds but lack intercellular effects between astrocytes and neurons. Astrocytes in coculture have a more ramified morphology (Martinez-Lozada et al., 2023) with fine structures contacting synapses (Araque et al., 1999). We hypothesized there would be differences in frequency or extent of astrocytic events in coculture with neurons, due to neurotransmitter- or neuronal activity-induced modulation of astrocytes (Gordon et al., 2005; Gordon et al., 2009; Perea & Araque, 2005; Porter & McCarthy, 1996).

Comparison of max projections of representative recordings of astrocyte monocultures and astrocyte-neuron cocultures shows a difference in morphology (Fig. 3A). Astrocytes without neurons are blocky, while astrocytes in cocultures form more processes and fine branches, which could reflect the presence of synapses. Spontaneous fluorescence changes in cocultured astrocytes were strikingly larger than those of astrocytes in monocultures (Fig. 3B). Furthermore, these events had a higher mean fluorescence (Fig. 3C; *p*<0.001) and event amplitude (Fig. 3D; β=2.94, 95%-CI: 2.75, 3.14, *p*<0.001). In raster plots, a higher density of events is visible in cocultures (Fig. 3E), due to a higher event frequency (Fig. 3F; *p*<0.001). No difference in overall event duration (Fig. 3G) was observed. However, there were significantly fewer fast and slow events (0-2 seconds: *p*=0.0038; >6 seconds: *p*=2.73×10^-4^), and more medium duration events (2-6 seconds: *p*<0.001) in cocultures compared to monocultures. The overall event area was lower in cocultured astrocytes (Fig. 3H; β=0.68, 95%-CI: 0.54, 0.85, *p*<0.001) due to more wavelet events (*p*=0.0063), and fewer waves (*p*<0.001). This indicates that the presence of neurons increases astrocytic calcium signaling frequency and amplitude of events, and causes a shift towards smaller-sized events, suggesting an increase in microdomain activity, potentially at perisynaptic sites. Statistics are summarized in Figure 3I and Table 3.

**Figure 3:**
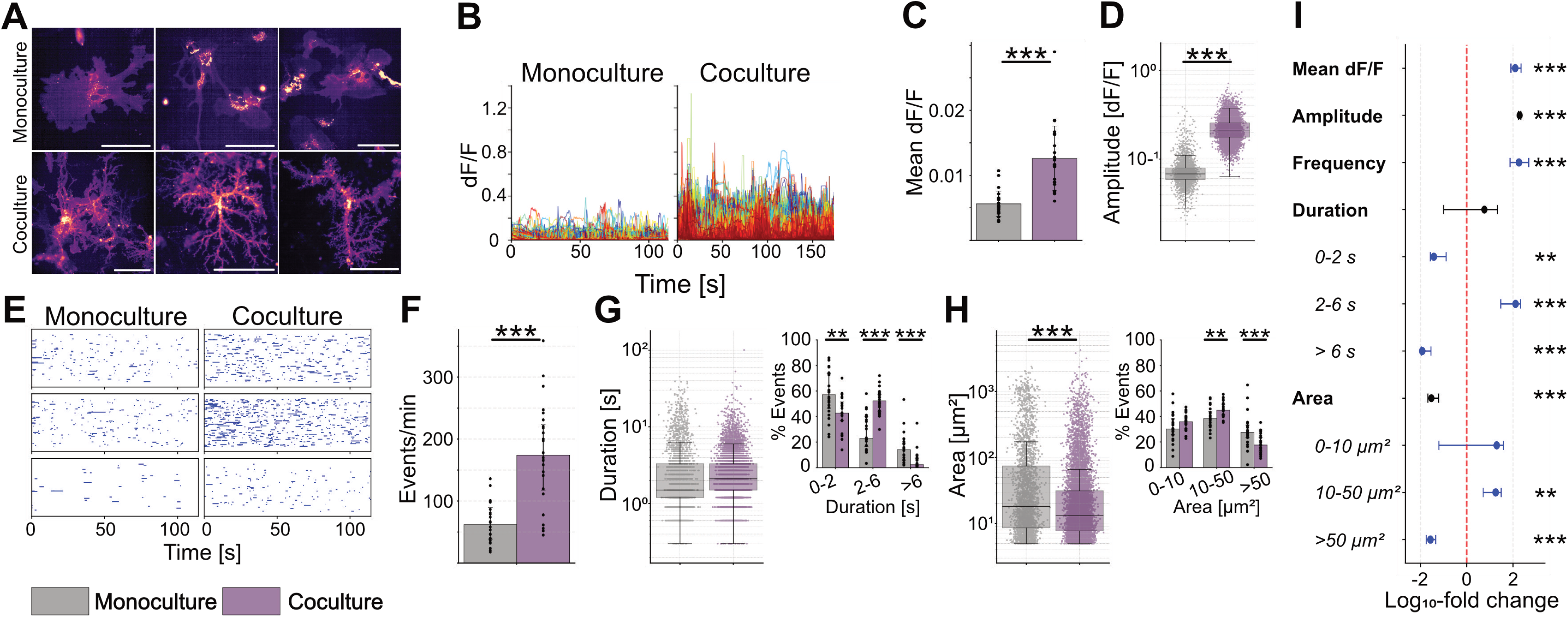
Baseline activity differs between mouse astrocytes in monoculture and in coculture with neurons. **(A)** Representative images of mouse astrocytes in monoculture or in coculture with neurons, transduced with AAV5 GFAP promoter-driven membrane-targeted GCaMP6f. **(B)** Individual event dF/F traces from monoculture and coculture recordings. **(C)** Mean fluorescence per sample from event traces, displayed per sample (dots) and group means (bars) ± SD. **(D)** Event amplitude; each dot is an individual event. Boxplots indicate median (center line), interquartile ranges (25^th^-75^th^ percentile), and whiskers extending to 1.5×IQR. **(E)** Raster plots showing occurrence of individual astrocytic events as blue horizontal lines, with length corresponding to event duration. **(F)** Event frequency, displayed per sample (dots) and group median (bars) ± IQR. **(G)** Event duration; each dot is an individual event. Boxplots indicate median (center line), interquartile ranges (25^th^-75^th^ percentile), and whiskers extending to 1.5×IQR. Inset (right) shows percent of events per duration subclass per sample (dots) and group median (bars) ± IQR. **(H)** Event area; each dot is an individual event. Boxplots indicate median (center line), interquartile ranges (25^th^-75^th^ percentile), and whiskers extending to 1.5×IQR. Inset (right) shows percent of events per area subclass per sample (dots) and group median (bars) ± IQR. **(I)** Forest plot summarizing differences per parameter as percent change on a log_10_-scale. Amplitude, duration, and area show effect size and confidence intervals derived from linear models in black. Mean dF/F, frequency, and duration and area subclasses, show per-sample percent change across all samples as median with interquartile ranges (25^th^-75^th^ percentile) in blue (Mann-Whitney-U-test). n = 24 each for astrocyte monoculture and coculture recordings; * p > 0.05, ** p > 0.01, *** p > 0.001. Scale bar is 100 µm.

**Table 3:** Descriptive statistics comparison of mouse astrocytes in monoculture & coculture with neurons.

|  | Mouse astrocyte monoculture | Mouse astrocyte-neuron coculture | % Change | p-value |
| --- | --- | --- | --- | --- |
| Frequency (events/min); median ± IQR; MWU | 61.94 (38.58–89.50) | 174.02 (117.59–223) | 180.93% | 6.01E-06 |
| Mean dF/F; mean ± SD; MWU | 0.01 ± 0.00 | 0.01 ± 0.00 | 124.18% | 6.97E-08 |
| Amplitude; median ± IQR; LMM | 0.07 (0.06–0.08) | 0.21 (0.18–0.26) | 193.56% | 1.45E-222 |
| Duration; median ± IQR; LMM | 1.50 (1.20–3.30) | 2.10 (1.50–3.30) | 4.87% | 0.521568 |
| Duration class 0–2 s; median ± IQR; MWU | 57.12 (46.36–73.76) | 42.67 (36.97–51.43) | -25.30% | 0.003767 |
| Duration class 2–6 s; median ± IQR; MWU | 22.85 (16.49–39.88) | 52.28 (44.20–57.14) | 128.78% | 4.62E-07 |
| Duration class >6 s; median ± IQR; MWU | 14.13 (5.41–19.30) | 2.45 (1.45–6.94) | -82.65% | 0.000273 |
| Area; median ± IQR; LMM | 18.20 (8.62–74.11) | 13.14 (7.76–31.24) | -32.42% | 0.000734 |
| Area class 0–10 μm<sup>2</sup>; median ± IQR; MWU | 30.23 (25.83–38.43) | 35.94 (30.38–39.61) | 18.90% | 0.077904 |
| Area class 10–50 μm<sup>2</sup>; median ± IQR; MWU | 38.42 (33.12–43.84) | 44.94 (41.20–49.74) | 16.97% | 0.006293 |
| Area class >50 μm<sup>2</sup>; median ± IQR; MWU | 27.59 (23.50–33.32) | 17.66 (12.92–23.72) | -35.97% | 0.000901 |

We then tested the effect of ATP and CPA on astrocytes in astrocyte-neuron cocultures. We saw a strong response to ATP (Fig. 4A, Movie 3), similar to astrocytes in monoculture. Fluorescent traces were elevated in height and width (Fig. 4B), and mean fluorescence was increased (Fig. 4C) within 30 seconds of addition of ATP (*p*<0.001). In contrast to monocultures, we did not observe an increase in the amplitude of individual events (Fig. 4D), despite the increase in mean fluorescence. This could be caused by more ATP-induced low amplitude events. Raster plots (Fig. 4E) appeared denser after ATP application, but unlike in astrocyte monocultures, this effect was not due to frequency, which was unchanged (Fig. 4F), but rather to increased event duration (Fig. 4G; β=1.95, 95%-CI: 1.59, 2.4, *p*<0.001), caused by a decrease in short 0-2 second events (*p*<0.001), and increase in long >6 second events (*p*<0.001). Event area, which was unchanged by ATP in monocultures, was increased by ATP in astrocytes in cocultures (Fig. 4H; β=1.75, 95%-CI: 1.28, 2.38, *p*<0.001), due to a decrease in wavelets (*p*=0.0068) and an increase in waves (*p*=0.021). Statistics are summarized in Figure 4I and Table 4. Thus, ATP increases astrocytic calcium signaling in both monocultures and cocultures, but the parameters affected (event amplitude, frequency, and area) are different.

**Figure 4:**
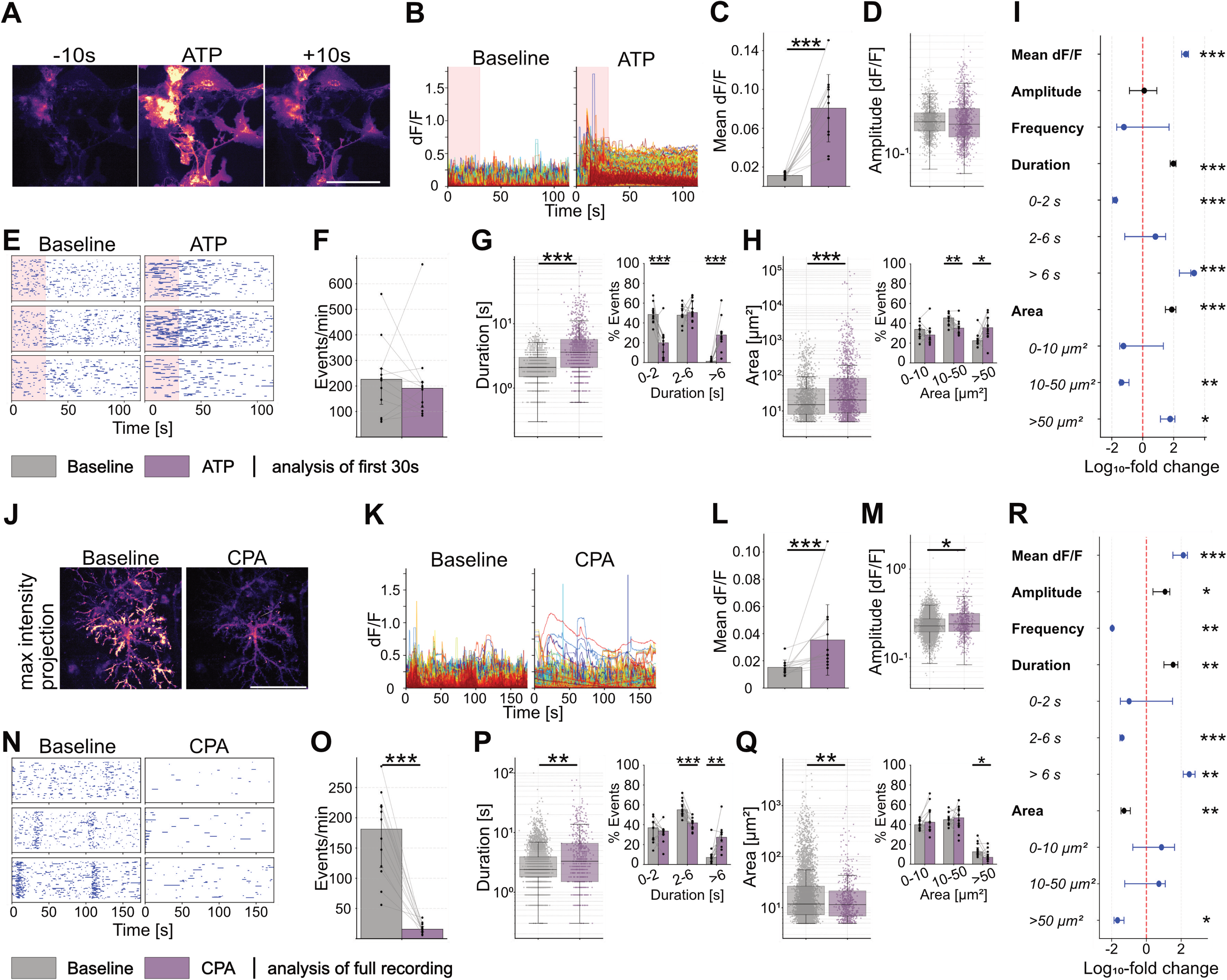
ATP increases, and CPA decreases calcium activity in mouse astrocytes in astrocyte-neuron cocultures. **(A)** Time-lapse images of a typical response induced by 50 µM ATP in mouse astrocytes in astrocyte-neuron cocultures transduced with AAV5 GFAP promoter-driven membrane-targeted GCaMP6f. **(B)** Individual event dF/F traces from baseline and treatment recordings. Light red shade indicates 30 second time window used for analysis. **(C)** Mean fluorescence per sample from event traces, displayed as paired data (dots) and group means (bars) ± SD. **(D)** Event amplitude; each dot is an individual event. Boxplots indicate median (center line), interquartile ranges (25^th^-75^th^ percentile), and whiskers extending to 1.5×IQR. **(E)** Raster plots showing occurrence of individual astrocytic events as blue horizontal lines, with length corresponding to event duration. **(F)** Event frequency, displayed as paired regions (dots) and group median (bars) ± IQR. **(G)** Event duration; each dot is an individual event. Boxplots indicate median (center line), interquartile ranges (25^th^-75^th^ percentile), and whiskers extending to 1.5×IQR. Inset (right) shows percent of events per duration subclass paired per-sample (dots) and group median (bars) ± IQR. **(H)** Event area; each dot is an individual event. Boxplots indicate median (center line), interquartile ranges (25^th^-75^th^ percentile), and whiskers extending to 1.5×IQR. Inset (right) shows percent of events per area subclass paired per-sample (dots) and group median (bars) ± IQR. **(I)** Forest plot summarizing treatment effects per parameter as percent change on a log_10_-scale. Amplitude, duration, and area show effect size and confidence intervals derived from linear mixed-effect models in black. Mean dF/F, frequency, and duration and area subclasses, show per-sample paired percent change across all samples as median with interquartile ranges (25^th^-75^th^ percentile) in blue (Wilcoxon signed-rank test). **(J)** Representative max intensity projections, **(K)** event dF/F traces, **(L)** mean fluorescence, **(M)** event amplitude, **(N)** raster plots, **(O)** event frequency, **(P)** event duration and sub-classes (inset), **(Q)** event area and sub-classes (inset), **(R)** and forest plot of mouse cocultured astrocytes transduced with AAV5 GFAP promoter-driven membrane-targeted GCaMP6f after 20 µM CPA treatment. n = 12 each for ATP and CPA baseline-treatment pairs; * p > 0.05, ** p > 0.01, *** p > 0.001. Scale bar is 100 µm.

**Table 4:** Descriptive statistics for effects of ATP and CPA in mouse astrocyte-neuron cocultures.

|  | Baseline | ATP | % Change | p-value |
| --- | --- | --- | --- | --- |
| Frequency (events/min); median ± IQR; WSR | 226.00 (128.00–268.00) | 191.00 (115.50–214.50) | -15.49% | 0.622070 |
| Mean dF/F; mean ± SD; WSR | 0.01 ± 0.00 | 0.08 ± 0.03 | 616.62% | 0.000488 |
| Amplitude; median ± IQR; LMM | 0.18 (0.15–0.21) | 0.17 (0.14–0.23) | 0.25% | 0.940419 |
| Duration; median ± IQR; LMM | 2.10 (1.50–3.00) | 3.60 (2.10–5.70) | 94.92% | 2.22E-10 |
| Duration class 0–2 s; median ± IQR; WSR | 48.53 (43.84–52.63) | 19.95 (4.84–25.59) | -58.89% | 0.000977 |
| Duration class 2–6 s; median ± IQR; WSR | 47.84 (46.57–54.70) | 50.53 (45.80–62.83) | 5.62% | 0.850098 |
| Duration class >6 s; median ± IQR; WSR | 1.28 (0.61–2.00) | 27.84 (23.05–30.94) | 2080.82% | 0.000488 |
| Area; median ± IQR; LMM | 15.08 (8.08–42.44) | 20.57 (8.83–84.39) | 74.65% | 0.000399 |
| Area class 0–10 $\mu\text{m}^2$; median ± IQR; WSR | 33.85 (26.62–37.74) | 28.20 (22.97–32.00) | -16.69% | 0.266113 |
| Area class 10–50 $\mu\text{m}^2$; median ± IQR; WSR | 45.46 (38.55–47.54) | 35.15 (29.81–37.21) | -22.69% | 0.006836 |
| Area class >50 $\mu\text{m}^2$; median ± IQR; WSR | 22.44 (19.84–25.10) | 35.60 (29.16–45.49) | 58.65% | 0.020996 |
|  | Baseline | CPA | % Change | p-value |
| Frequency (events/min); median ± IQR; WSR | 181.38 (117.59–218.53) | 15.69 (11.21–20.52) | -91.35% | 0.000488 |
| Mean dF/F; mean ± SD; WSR | 0.02 ± 0.01 | 0.04 ± 0.03 | 131.83% | 0.000488 |
| Amplitude; median ± IQR; LMM | 0.23 (0.20–0.28) | 0.24 (0.20–0.32) | 10.84% | 0.023536 |
| Duration; median ± IQR; LMM | 2.40 (1.80–3.90) | 3.30 (1.50–6.60) | 33.62% | 0.004894 |
| Duration class 0–2 s; median ± IQR; WSR | 36.89 (25.24–42.67) | 33.63 (30.14–34.81) | -8.82% | 0.569336 |
| Duration class 2–6 s; median ± IQR; WSR | 55.03 (50.98–60.70) | 41.87 (36.84–43.12) | -23.92% | 0.000488 |
| Duration class >6 s; median ± IQR; WSR | 7.00 (3.11–9.66) | 27.37 (22.59–32.49) | 291.17% | 0.002441 |
| Area; median ± IQR; LMM | 11.85 (7.32–26.50) | 11.53 (6.89–21.44) | -18.24% | 0.001518 |
| Area class 0–10 $\mu\text{m}^2$; median ± IQR; WSR | 39.84 (35.90–44.22) | 42.40 (34.82–54.55) | 6.41% | 0.203613 |
| Area class 10–50 $\mu\text{m}^2$; median ± IQR; WSR | 44.94 (40.74–48.67) | 46.85 (38.96–54.66) | 4.23% | 0.909668 |
| Area class >50 $\mu\text{m}^2$; median ± IQR; WSR | 12.52 (10.28–16.88) | 6.93 (4.85–10.34) | -44.68% | 0.026855 |

CPA induced a reliable decrease in fluorescence intensity of astrocytic calcium signaling in maximum intensity projections in astrocyte-neuron cocultures (Fig. 4J), with a few events that showed prolonged fluorescence increases (Fig. 4K) similar to astrocyte monocultures. However, unlike monocultures, CPA increased mean fluorescence (Fig. 4L; *p*<0.001) and event amplitude (Fig. 4M; β=1.11, 95%-CI: 1.01, 1.21, *p*=0.024), contrasting the decrease in amplitude seen in monocultures. Raster plots (Fig. 4N) indicated a pronounced decrease in overall activity, confirmed by a decrease in event frequency (Fig. 4O; *p*<0.001), similar to monocultures. The event duration was similarly increased (Fig. 4P; β=1.34, 95%-CI: 1.09, 1.64, *p*=0.0049), due to a reduction in 2-6 second events (*p*<0.001) and an increase in >6 second events (*p*=0.0024). Event area decreased (Fig. 4Q; β=0.82, 95%-CI: 0.72, 0.93, *p*=0.0015), also similar to astrocyte monocultures, due to a decrease in waves (*p*=0.027).

Although CPA caused an overall decrease in calcium activity in astrocytes in both monocultures and cocultures, some parameters differed, i.e., event amplitude was decreased in monocultures but increased in cocultures. Statistics are summarized in Figure 4R and Table 4.

### Glutamate and MPEP induced calcium responses in astrocytes in coculture

While astrocyte monocultures isolate astrocyte-specific effects of compounds, cocultures represent a more physiological system in which compounds may affect both cell types. We therefore tested glutamate and MPEP, which are known to affect both cell types, and measured astrocytic effects using our pipeline. Astrocytes express a variety of glutamate receptors (Dzamba et al., 2015; Lalo et al., 2011; Verkhratsky & Kirchhoff, 2007), and glutamate is known to increase calcium in astrocytes (Cai et al., 2000; James et al., 2011; Porter & McCarthy, 1996). In cocultures, glutamate triggered robust calcium responses in astrocytes (Fig. 5A), with a rapid elevation of fluorescence intensity in traces (Fig. 5B) and an increase in mean fluorescence (Fig. 5C; *p*<0.001). However, event amplitude was decreased by glutamate (β=0.90, 95%-CI: 0.84, 0.97, *p*=0.0082) (Fig. 5D), likely due to the appearance of more low-amplitude events. Raster plots (Fig. 5E) showed increases in event frequency (Fig. 5F; *p*<0.001) and duration (Fig. 5G; β=1.89, 95%-CI: 1.68, 2.12, *p*<0.001) due to a decrease in short 0-2 second events (*p*=6.10×10^-5^) and increase in longer events (2-6 seconds: *p*<0.001; >6 seconds: *p*<0.001). Event area was increased (Fig. 5H; β=1.57, 95%-CI: 1.24, 2.00, *p*<0.001), due to an increase in wave events (*p*=0.010). Statistics are summarized in Figure 5I and Table 5.

**Figure 5:**
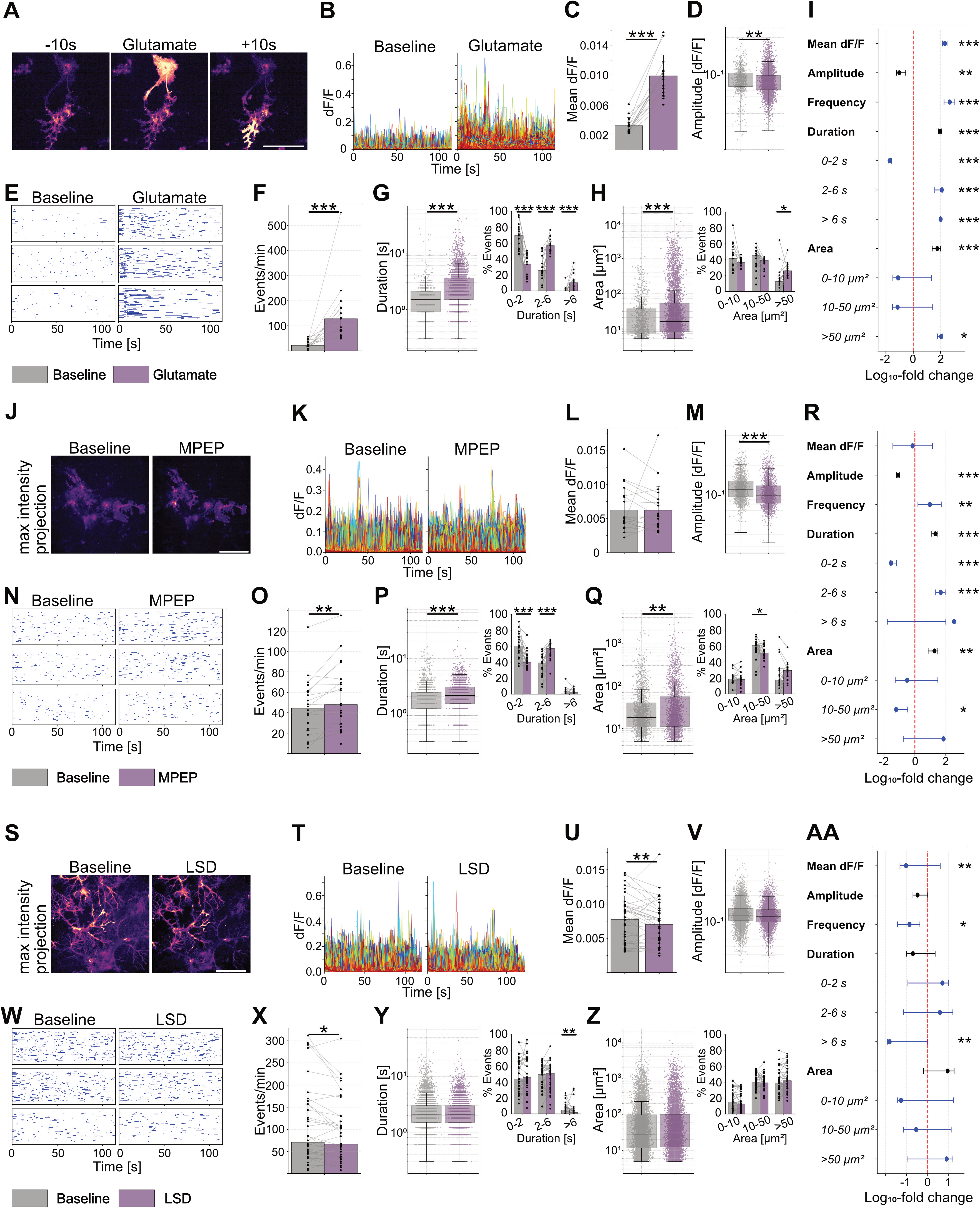
Glutamate, MPEP, and LSD elicit specific astrocyte calcium signals in astrocyte-neuron cocultures. **(A)** Time-lapse images of a typical response induced by 100 µM Glutamate in mouse astrocytes in astrocyte-neuron cocultures transduced with AAV5 GFAP promoter-driven membrane-targeted GCaMP6f. **(B)** Individual event dF/F traces from baseline and treatment recordings. **(C)** Mean fluorescence per sample from event traces, displayed as paired data (dots) and group means (bars) ± SD. **(D)** Event amplitude; each dot is an individual event. Boxplots indicate median (center line), interquartile ranges (25^th^-75^th^ percentile), and whiskers extending to 1.5×IQR. **(E)** Raster plots showing occurrence of individual astrocytic events as blue horizontal lines, with length corresponding to event duration. **(F)** Event frequency, displayed as paired regions (dots) and group median (bars) ± IQR. **(G)** Event duration; each dot is an individual event. Boxplots indicate median (center line), interquartile ranges (25^th^-75^th^ percentile), and whiskers extending to 1.5×IQR. Inset (right) shows percent of events per duration subclass paired per-sample (dots) and group median (bars) ± IQR. **(H)** Event area; each dot is an individual event. Boxplots indicate median (center line), interquartile ranges (25^th^-75^th^ percentile), and whiskers extending to 1.5×IQR. Inset (right) shows percent of events per area subclass paired per-sample (dots) and group median (bars) ± IQR. **(I)** Forest plot summarizing treatment effects per parameter as percent change on a log_10_-scale. Amplitude, duration, and area show effect size and confidence intervals derived from linear mixed-effect models in black. Mean dF/F, frequency, and duration and area subclasses, show per-sample paired percent change across all samples as median with interquartile ranges (25^th^-75^th^ percentile) in blue (Wilcoxon signed-rank test). **(J)** Representative max intensity projections, **(K)** event dF/F traces, **(L)** mean fluorescence, **(M)** event amplitude, **(N)** raster plots, **(O)** event frequency, **(P)** event duration and sub-classes (inset), **(Q)** event area and sub-classes (inset), **(R)** and forest plot of mouse cocultured astrocytes transduced with AAV5 GFAP promoter-driven membrane-targeted GCaMP6f after 50 µM MPEP treatment. **(S)** Representative max intensity projections, **(T)** event dF/F traces, **(U)** mean fluorescence, **(V)** event amplitude, **(W)** raster plots, **(X)** event frequency, **(Y)** event duration and sub-classes (inset), **(Z)** event area and sub-classes (inset), **(AA)** and forest plot of mouse cocultured astrocytes transduced with AAV5 GFAP promoter-driven membrane-targeted GCaMP6f one hour after 10 µM LSD treatment. n = 15 (Glutamate), 20 (MPEP), and 36 (LSD) baseline-treatment pairs; * p > 0.05, ** p > 0.01, *** p > 0.001. Scale bar is 100 µm.

**Table 5:**
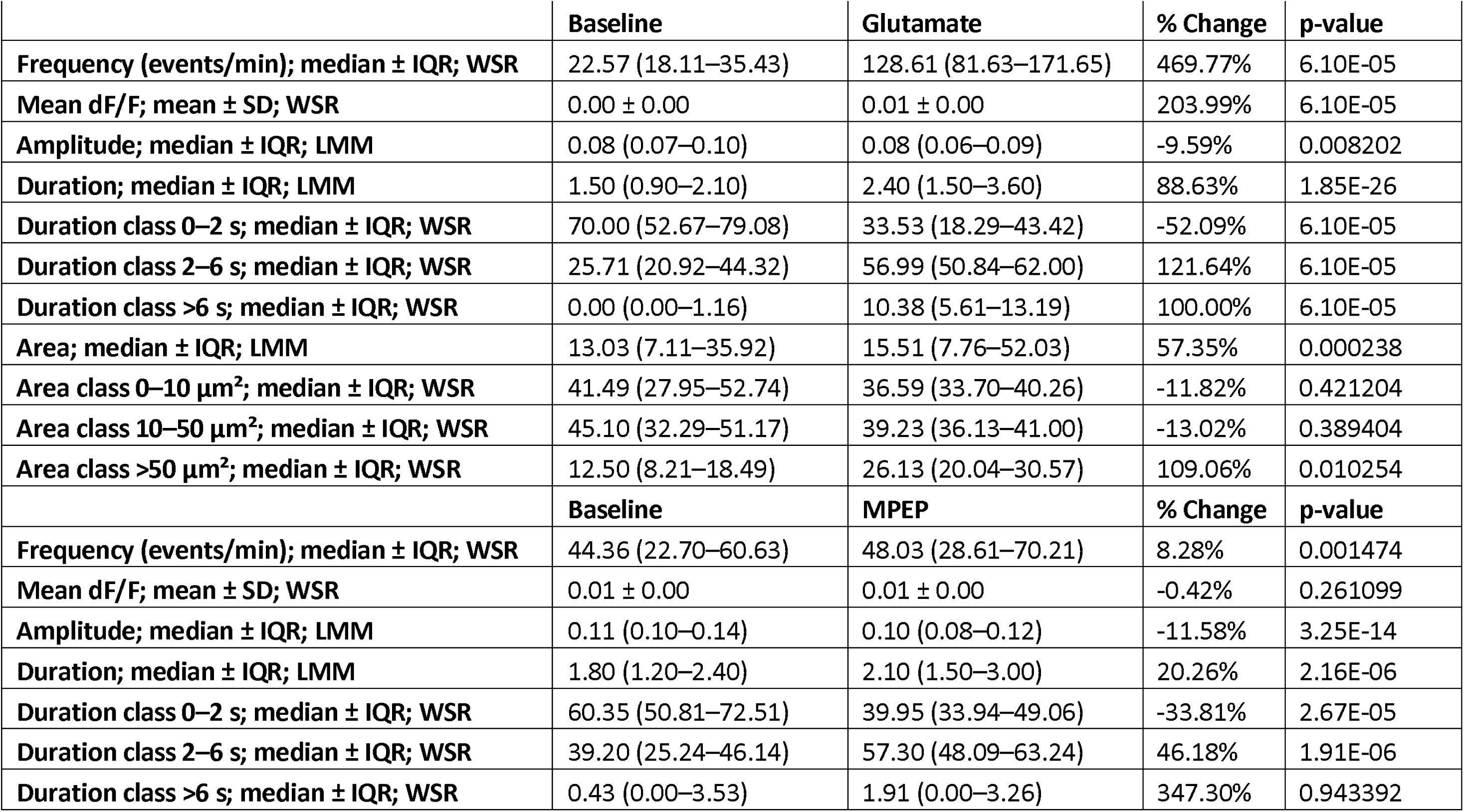
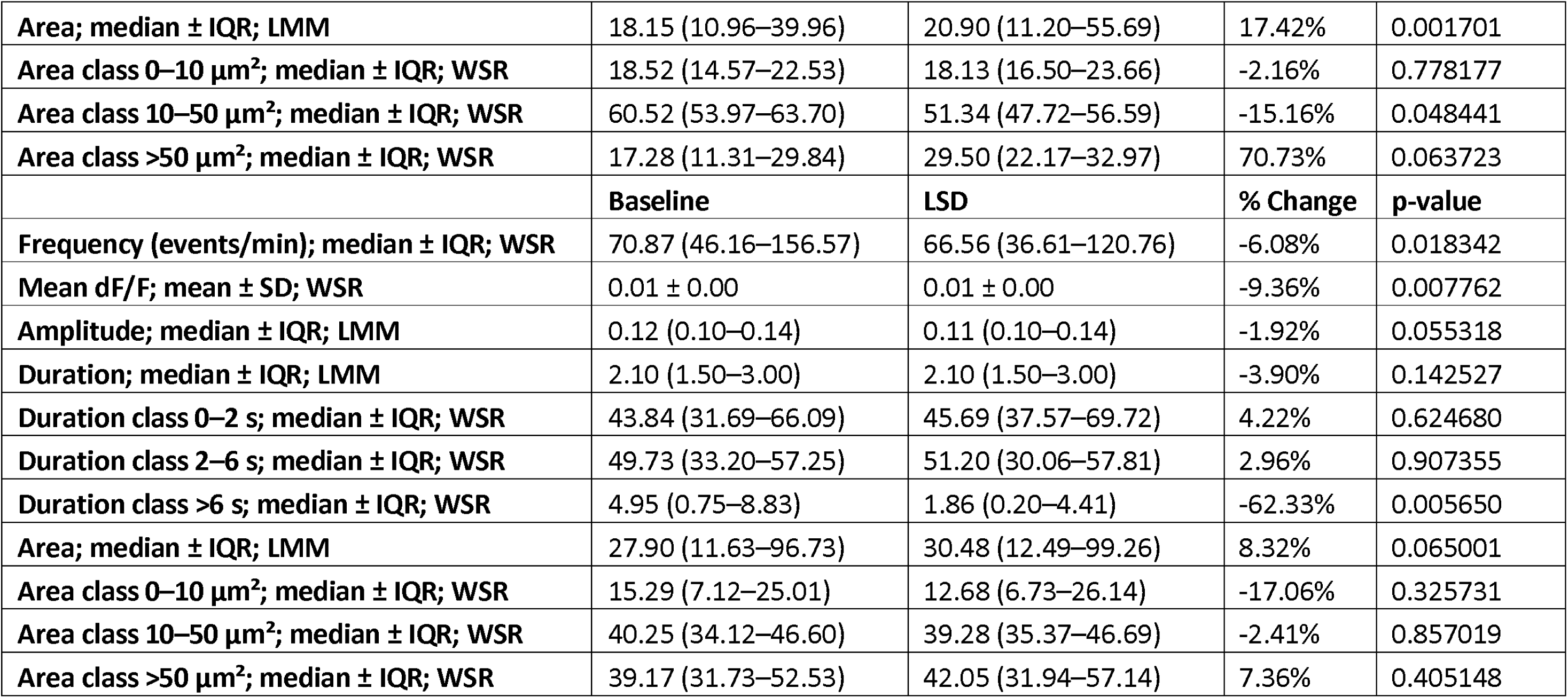
Descriptive statistics effects of glutamate, MPEP, and LSD in mouse astrocyte-neuron cocultures.

MPEP is an antagonist of the mGluR5 receptor, which is expressed in young astrocytes. If the receptor is present, we would expect a decrease in activity (Panatier et al., 2011). In tests of MPEP in cocultures, we found no change in maximum intensity projections of baseline and treated cells (Fig. 5J), fluorescent traces (Fig. 5K), or mean fluorescence (Fig. 5L). Event amplitude was decreased after application of MPEP (Fig. 5M; β=0.88, 95%-CI: 0.86, 0.91, *p*<0.001). Raster plots (Fig. 5N) indicated a slight increase in event frequency (Fig. 5O; *p*=0.0015) and duration (Fig. 5P; β=1.20, 95%-CI: 1.11, 1.30, *p*<0.001), due to a shift to slower events, with a decrease in 0-2 second events (*p*<0.001) and increase in 2-6 second events (*p*<0.001). Event area was also increased (Fig. 5Q; β=1.17, 95%-CI: 1.06, 1.3, *p*=0.0017), due to fewer wavelets (*p*=0.048). Statistics are summarized in Figure 5R and Table 5. Only one parameter (amplitude) showed the expected inhibitory effect seen in young astrocytes (Panatier et al., 2011), while the majority of event parameters (frequency, duration, and area) were increased by MPEP. This could indicate that our astrocytes are mature (Cai et al., 2000; Sun et al., 2013).

### LSD decreases event frequency and duration, and increases event area in mouse astrocytes

We next tested lysergic acid diethylamide (LSD), a psychoactive drug that targets serotonin receptors and has not previously been tested for effects on astrocyte calcium signaling. Following treatment of mouse astrocyte-neuron cocultures with LSD for one hour, maximum intensity projections (Fig. 5S) and fluorescent traces (Fig. 5T) appeared to decrease, reflected by a decrease in mean fluorescence (Fig. 5U; *p*=0.0078). Raster plots also showed a slight decrease in event density (Fig. 5W) caused by a decrease in event frequency (Fig. 5X; *p*=0.018). While overall event duration remained unchanged (Fig. 5Y), we found a reduction in slow >6 second events (*p*=0.0057). Event area (Fig. 5Z) was unchanged. Statistics are summarized in Figure 5AA and Table 5. Using our pipeline, we report that LSD reduces astrocytic calcium signaling in mouse astrocytes.

### Dual-color calcium imaging reveals effects of gabazine on both neurons and astrocytes

To assess astrocytic effects of compounds relative to neuronal firing, we expanded the pipeline using a combination of green (GCaMP) and red (RCaMPs, jRGECO) fluorescent calcium indicators, expressed in neurons (using the synapsin promoter) or astrocytes (using the GFAP promoter). We used membrane-targeted GFAP promoter-driven GCaMP6f in combination with synapsin promoter-driven RCaMP1a, or membrane-targeted GFAP promoter-driven jRGECO together with synapsin promoter-driven GCaMP6f, and data from both combinations were combined.

We first tested gabazine, a GABA_A_ receptor antagonist (Ueno et al., 1997), which induces disinhibition and synchronous firing of neurons in culture (Pegoraro et al., 2010). Astrocytes have been shown to express GABA receptors themselves (Höft et al., 2014) and respond to GABA receptor agonists (Fraser et al., 1995; Meier et al., 2008; Nilsson et al., 1993). Dual-color imaging yielded specific signals for neurons or astrocytes (Fig. 6A). Astrocyte event traces (Fig. 6B) showed a trend toward decreased mean fluorescence (Fig. 6C), and astrocyte event amplitude was decreased by gabazine treatment (Fig. 6D; β=0.91, 95%-CI: 0.86, 0.96, *p*<0.001). Raster plots (Fig. 6E) indicated a decrease in astrocyte event frequency (*p*<0.001) and increase in synchronous neuronal firing frequency (*p*=0.032) (Fig. 6F). Astrocyte event duration (Fig. 6G) remained unchanged, but event area was reduced (Fig. 6H; β=0.83, 95%-CI: 0.72, 0.94, *p*=0.0049) due to a reduction in the size of calcium waves (*p*=0.032). Statistics are summarized in Figure 6I and Table 6. This approach can be used to simultaneously test the effects of compounds on neurons and astrocytes.

**Figure 6:**
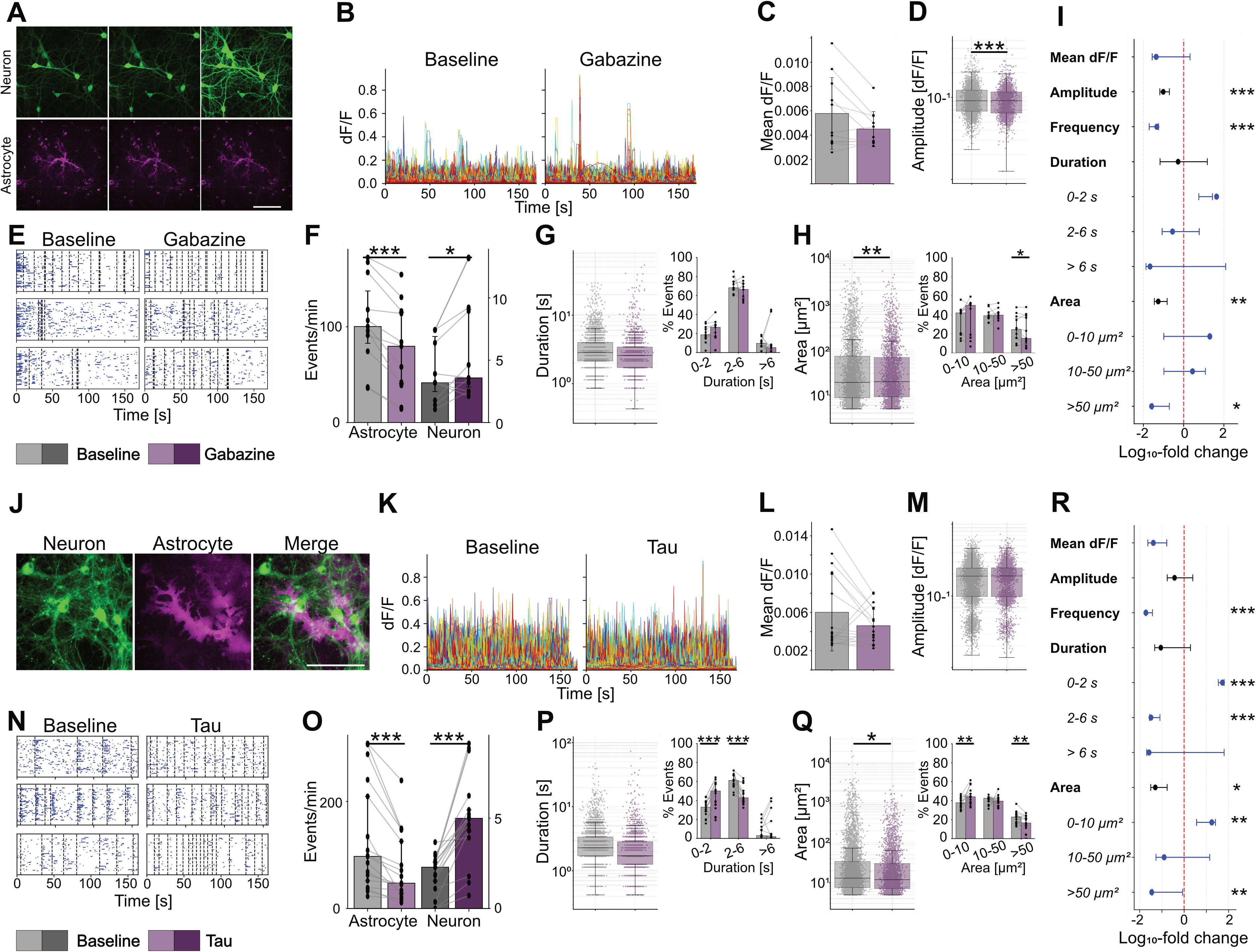
Dual-color imaging reveals differential effects of gabazine and Tau on astrocytes and neurons. **(A)** Time-lapse images of a typical response induced by 10 µM gabazine in mouse astrocyte-neuron cocultures. Astrocytes (magenta) transduced with AAV5 GFAP promoter-driven membrane-targeted GCaMP6f or AAV5 GFAP promoter-driven membrane-targeted jRGECO, neurons (green) transduced with AAV1/2 synapsin promoter driven jRCaMP1a or AAV1 synapsin promoter driven GCaMP6f. **(B)** Individual astrocyte event dF/F traces from baseline and treatment recordings. **(C)** Mean fluorescence per sample from event traces, displayed as paired data (dots) and group means (bars) ± SD. **(D)** Event amplitude; each dot is an individual event. Boxplots indicate median (center line), interquartile ranges (25^th^-75^th^ percentile), and whiskers extending to 1.5×IQR. **(E)** Raster plots showing occurrence of individual astrocytic events as blue horizontal lines, with length corresponding to event duration, and occurrence of neuron events as dashed vertical lines. **(F)** Event frequency, displayed as paired regions (dots) and group median (bars) ± IQR for astrocytes and neurons. **(G)** Event duration; each dot is an individual event. Boxplots indicate median (center line), interquartile ranges (25^th^-75^th^ percentile), and whiskers extending to 1.5×IQR. Inset (right) shows percent of events per duration subclass paired per-sample (dots) and group median (bars) ± IQR. **(H)** Event area; each dot is an individual event. Boxplots indicate median (center line), interquartile ranges (25^th^-75^th^ percentile), and whiskers extending to 1.5×IQR. Inset (right) shows percent of events per area subclass paired per-sample (dots) and group median (bars) ± IQR. **(I)** Forest plot summarizing treatment effects on astrocytes per parameter as percent change on a log_10_-scale. Amplitude, duration, and area show effect size and confidence intervals derived from linear mixed-effect models in black. Mean dF/F, frequency, and duration and area subclasses, show per-sample paired percent change across all samples as median with interquartile ranges (25^th^-75^th^ percentile) in blue (Wilcoxon signed-rank test). **(J)** Representative max intensity projections, **(K)** event dF/F traces, **(L)** mean fluorescence, **(M)** event amplitude, **(N)** raster plots, **(O)** event frequency, **(P)** event duration and sub-classes (inset), **(Q)** event area and sub-classes (inset), **(R)** and forest plot of mouse astrocyte-neuron cocultures four hours after treatment with 25 nM human Tau oligomers; Astrocytes (magenta) transduced with AAV5 GFAP promoter-driven membrane-targeted GCaMP6f or AAV5 GFAP promoter-driven membrane-targeted jRGECO, neurons (green) transduced with AAV1/2 synapsin promoter driven jRCaMP1a or AAV1 synapsin promoter driven GCaMP6f. n = 11 (gabazine) and 17 (Tau) baseline-treatment pairs; * p > 0.05, ** p > 0.01, *** p > 0.001. Scale bar is 100 µm.

**Table 6:**
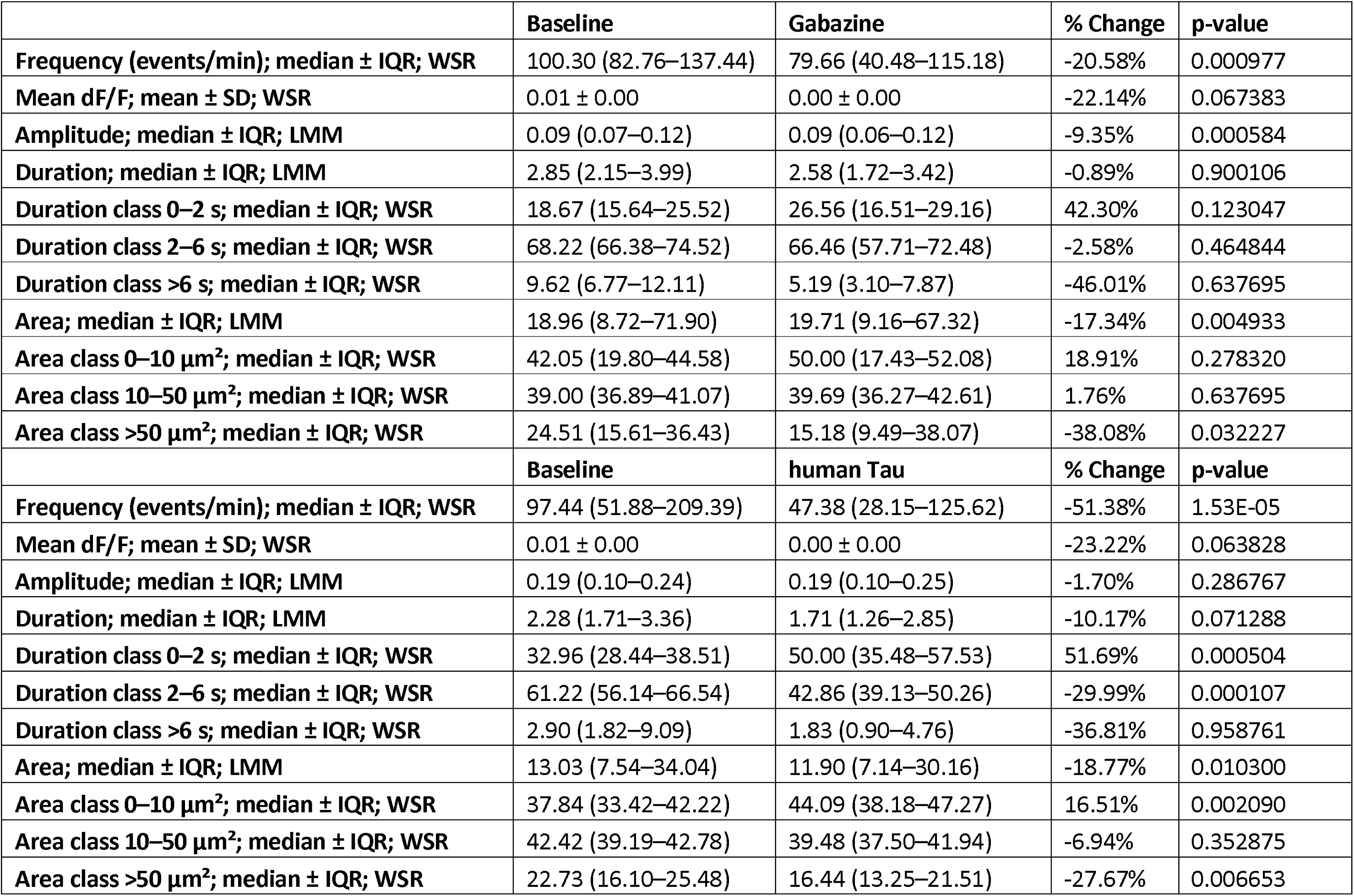
Descriptive statistics effects of gabazine and human Tau in mouse astrocyte-neuron cocultures.

### Astrocytes and neurons are differentially affected by extracellular Tau

To test the pipeline in a simplified disease model, we determined the effect of extracellular Tau oligomers, which are thought to contribute to the degeneration of neurons in Alzheimer’s disease (Busche & Hyman, 2020) on astrocytic and neuronal calcium signaling. We applied human recombinant Tau oligomers to cocultures for four hours and simultaneously recorded calcium activity from astrocytes and neurons expressing membrane-targeted GFAP promoter-driven GCaMP6f in combination with synapsin promoter-driven RCaMP1a, or membrane-targeted GFAP promoter-driven jRGECO together with synapsin promoter-driven GCaMP6f (Fig. 6J, Movie 4). We did not observe differences in fluorescence traces (Fig. 6K), mean fluorescence intensity (Fig. 6L), or event amplitude (Fig. 6M) of astrocytes upon treatment. However, raster plots (Fig. 6N) indicated a decrease in astrocyte event frequency (*p*<0.001) and an increase in neuronal event frequency (*p*<0.001) (Fig. 6O). The overall astrocytic event duration was unchanged (Fig. 6P), but 0-2 second events were increased (*p*<0.001) and 2-6 second events were decreased (*p*<0.001), indicating a shift to faster events induced by Tau oligomers. Astrocyte event area decreased (Fig. 6Q; β=0.81, 95%-CI: 0.69, 0.95, *p*=0.010), with an increase in microdomain activity (*p*=0.0021) and a decrease in wave events (*p*=0.0067). Statistics are summarized in Figure 6R and Table 6. Our pipeline was thus able to detect changes in both astrocytes and neurons after incubation with Tau oligomers.

### Mouse and human iPSC-derived astrocytes show differences in calcium signaling

Because mouse models can have low predictive value for toxicity and efficacy of drugs in human trials (Atkins et al., 2020; Bailey et al., 2014; Marshall et al., 2023; Pound & Ritskes-Hoitinga, 2018; van Meer et al., 2012; Van Norman, 2019), we extended our pipeline to human astrocytes derived from two iPSC lines, to explore similarities and differences between mouse and human astrocytic calcium responses. We used BIHi005-A-1E (iAstrocytes) expressing NFiB and Sox9 under the inducible Tet-promoter (Breuer et al., 2026) and ioAstrocytes from bit.bio, similarly induced by forward programming but derived from a different donor background. Human astrocytes were examined in monoculture to enable detection of astrocytic calcium signals using the bath-applied calcium dye Cal520-AM, which is comparable in kinetics to GCaMP6f, without interference from neuronal activity, and were examined in baseline conditions.

Raster plots of baseline activity showed a higher density of events in iAstrocytes and ioAstrocytes compared to mouse astrocyte monocultures (Fig. 7A). Event frequency was similar in iAstrocytes and mouse astrocytes, but 2-fold higher in ioAstrocytes (Fig. 7B; ioAstrocyte vs. mouse: *p*<0.001; ioAstrocytes vs. iAstrocytes: *p*<0.001). Mean fluorescence was considerably higher in both iAstrocytes and ioAstrocytes than in mouse astrocytes, and also higher in ioAstrocytes compared to iAstrocytes (Fig. 7C; iAstrocytes vs. mouse: *p*<0.001, ioAstrocyte vs. mouse: *p*<0.001; ioAstrocytes vs. iAstrocytes: *p*=0.014). Event amplitude was lowest in mouse astrocytes, higher in iAstrocytes, and highest in ioAstrocytes (Fig. 7D, iAstrocytes vs. mouse: *p*<0.001, ioAstrocyte vs. mouse: *p*<0.001; ioAstrocytes vs. iAstrocytes: *p*<0.001). Event duration was shortest in mouse astrocytes, higher in iAstrocytes, and highest in ioAstrocytes (Fig. 7E; iAstrocytes vs. mouse: *p*<0.001, ioAstrocyte vs. mouse: *p*<0.001; ioAstrocytes vs. iAstrocytes: p=0.002). Classification of event durations revealed the near absence of fast 0-2 second events in both human astrocyte cultures; more than 50% of events were slow, >6 second events. In contrast, mouse astrocytes showed a low proportion of slow but a high proportion of fast events (0-2 s events: iAstrocytes vs. mouse: *p*<0.001, ioAstrocyte vs. mouse: *p*<0.001; ioAstrocytes vs. iAstrocytes: *p*<0.001; >6 s events: iAstrocytes vs. mouse: *p*<0.001, ioAstrocyte vs. mouse: *p*<0.001; ioAstrocytes vs. iAstrocytes: *p*<0.001). Mouse astrocytes displayed the smallest event areas, with larger events in iAstrocytes, and the largest events in ioAstrocytes (Fig. 7F; iAstrocytes vs. mouse: *p*<0.001, ioAstrocyte vs. mouse: *p*<0.001). This was due to a lower proportion of microdomain events and a higher proportion of waves in human astrocytes (microdomains: ioAstrocyte vs. mouse: *p*<0.001; ioAstrocytes vs. iAstrocytes: *p*<0.001; wavelets: iAstrocytes vs. mouse: *p*<0.001, ioAstrocyte vs. mouse: *p*<0.001; waves: iAstrocytes vs. mouse: *p*<0.001, ioAstrocyte vs. mouse: *p*<0.001; ioAstrocytes vs. iAstrocytes: *p*<0.001). However, some differences between mouse and human astrocytes could be caused by differences in the calcium indicator used. For example, GCaMP6f labels fine processes better than bath-applied dyes like Cal520 (Reeves et al., 2011; Shigetomi et al., 2013), which could explain the greater proportion of small events in mouse astrocytes labelled with GCaMP6f, compared to human astrocytes labelled with Cal520. Descriptive statistics can be found in Table 7. In summary, human astrocytes showed more events that were larger and slower than in mouse astrocytes.

**Figure 7:**
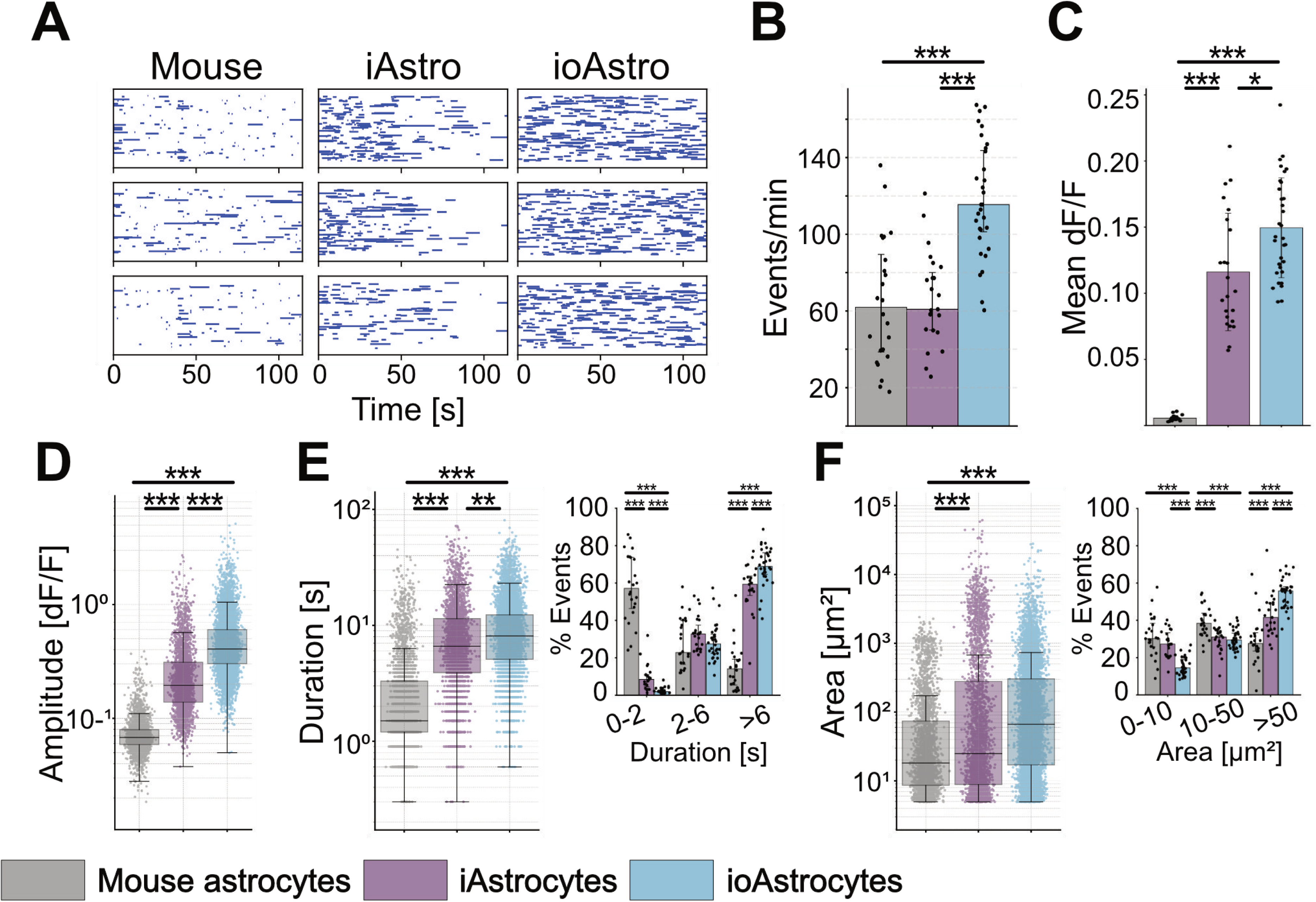
Comparison of baseline recordings from mouse astrocytes in monoculture, human iAstrocytes, and human ioAstrocytes. **(A)** Raster plots showing occurrence of individual astrocytic events as blue horizontal lines, with length corresponding to event duration. **(B)** Event frequency displayed per sample (dots) and group median (bars) ± IQR. **(C)** Mean fluorescence per sample from event traces, displayed per sampe (dots) and group means (bars) ± SD. **(D)** Event amplitude; each dot is an individual event. Boxplots indicate median (center line), interquartile ranges (25^th^-75^th^ percentile), and whiskers extending to 1.5×IQR. **(E)** Event duration; each dot is an individual event. Boxplots indicate median (center line), interquartile ranges (25^th^-75^th^ percentile), and whiskers extending to 1.5×IQR. Inset (right) shows percent of events per duration subclass per sample (dots) and group median (bars) ± IQR. **(F)** Event area; each dot is an individual event. Boxplots indicate median (center line), interquartile ranges (25^th^-75^th^ percentile), and whiskers extending to 1.5×IQR. Inset (right) shows percent of events per area subclass per sample (dots) and group median (bars) ± IQR. Analysis with linear mixed-effects model or Kruskal-Wallis test with Bonferroni FWER adjustments. n = 24 (mouse astrocytes), 24 (iAstrocytes), and 32 (ioAstrocytes) baseline recordings; * p > 0.05, ** p > 0.01, *** p > 0.001.

**Table 7:**
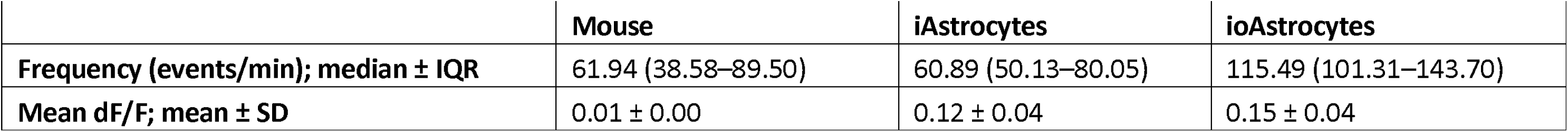
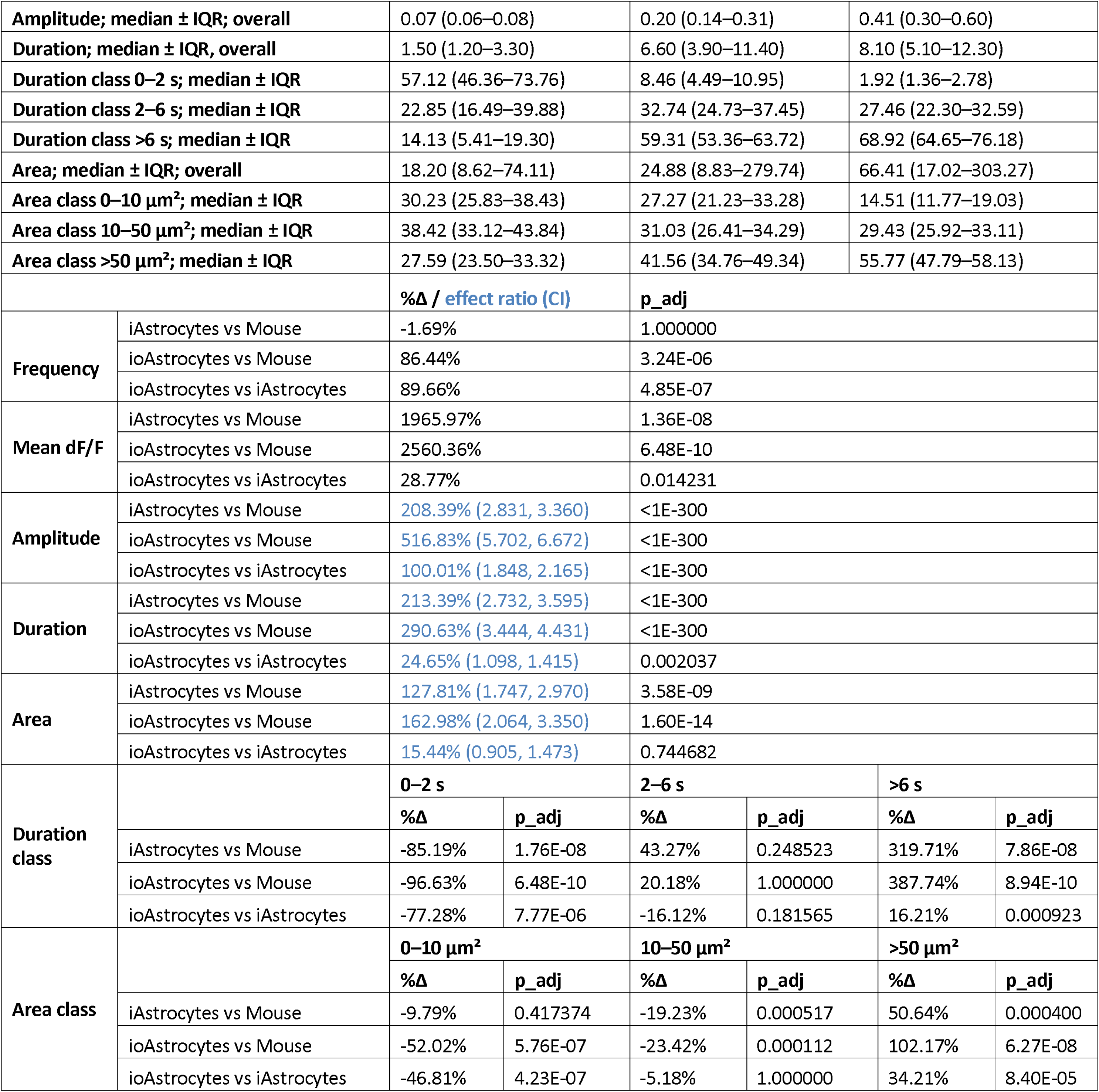
Descriptive statistics comparison of mouse astrocytes and human iAstrocytes and ioAstrocytes.

### Human iPSC-derived astrocyte calcium signaling is activated by ATP

We then examined the response of human astrocytes to ATP. ATP induced a rapid and robust activation of iAstrocytes (Fig. 8A, Movie 5), similar to mouse astrocytes and consistent with previous reports on primary human astrocytes (Oberheim et al., 2009; Zhang et al., 2016). Fluorescence traces showed a transient activation during the first 30 seconds (Fig. 8B) and an increase in mean fluorescence (Fig. 8C; *p*=0.018), but with a lower magnitude than mouse astrocytes. Individual regions did not always increase, and some even decreased in intensity, indicating more heterogeneity in human astrocytes. Event amplitude was increased by ATP (Fig. 8D; β=1.21, 95%-CI: 1.05, 1.41, *p*=0.010), similar to mouse astrocytes. Raster plots (Fig. 8E) showed a decrease in event frequency (Fig. 8F; *p*<0.001) in contrast to mouse astrocytes, and an increase in event duration (Fig. 8G) (β=1.32, 95%-CI: 1.13, 1.55, *p*<0.001) similar to mouse astrocytes. In iAstrocytes, the decrease in duration was due to fewer 2-6 second events (*p*<0.001) and more >6 second events (*p*<0.001), while in mouse astrocytes, it was due to fewer 0-2 second events. Unlike mouse astrocytes in monoculture, but similar to mouse coculture astrocytes, event area was increased in iAstrocytes after ATP application (Fig. 8H; β=1.90, 95%-CI: 1.23, 2.93, *p*=0.0035), due to a decrease in wavelets area (*p*=0.0031) and an increase in waves (*p*=0.0067). Statistics are summarized in Figure 8I and Table 8.

**Figure 8:**
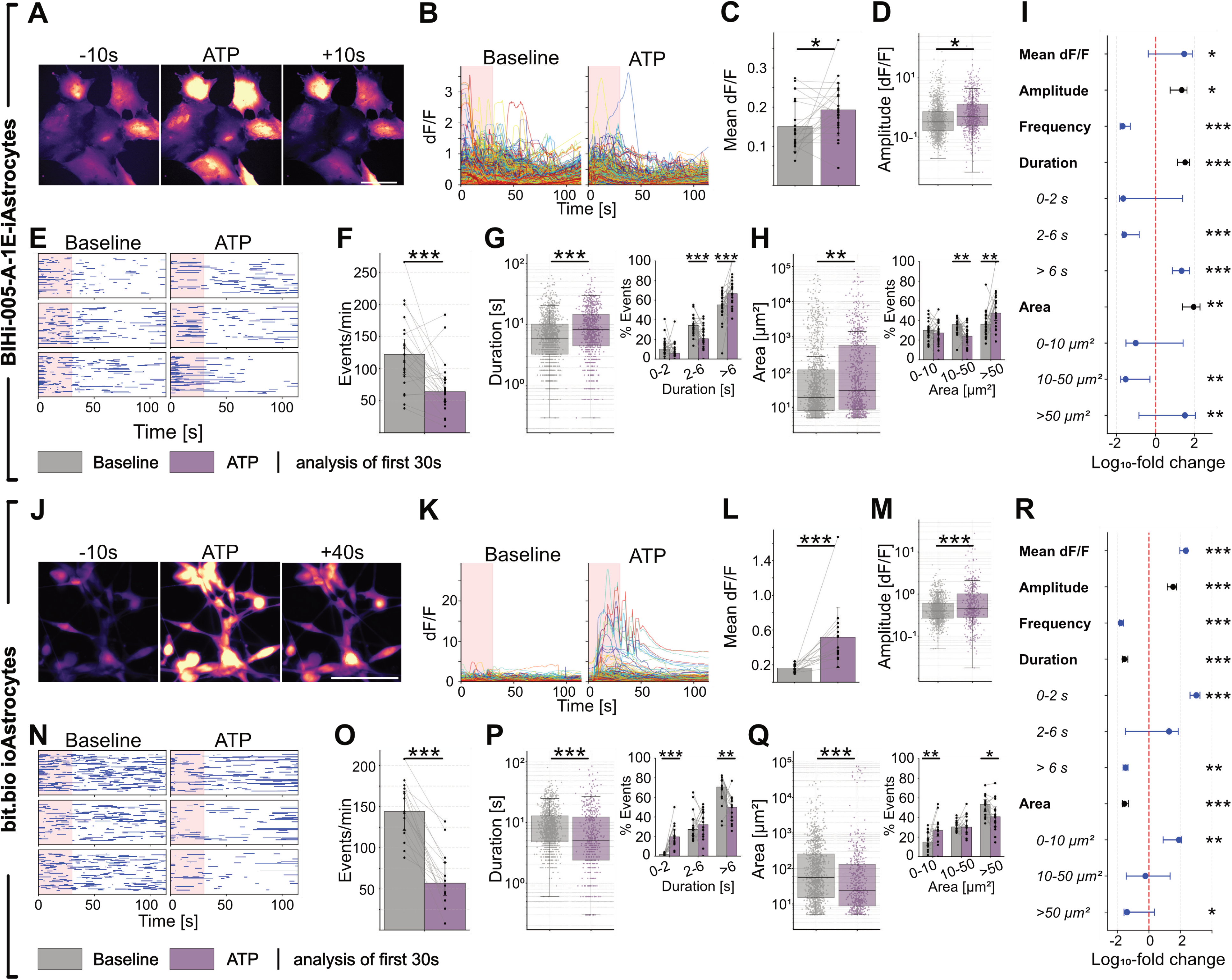
ATP increases calcium activity in human iAstrocytes and ioAstrocytes. **(A)** Time-lapse images of a typical response induced by 50 µM ATP in human iAstrocytes loaded with 2 µM Cal520-AM calcium dye. **(B)** Individual event dF/F traces from baseline and treatment recordings. Light red shade indicates 30 second time window used for analysis. **(C)** Mean fluorescence per sample from event traces, displayed as paired data (dots) and group means (bars) ± SD. **(D)** Event amplitude; each dot is an individual event. Boxplots indicate median (center line), interquartile ranges (25^th^-75^th^ percentile), and whiskers extending to 1.5×IQR. **(E)** Raster plots showing occurrence of individual astrocytic events as blue horizontal lines, with length corresponding to event duration. **(F)** Event frequency, displayed as paired regions (dots) and group median (bars) ± IQR. **(G)** Event duration; each dot is an individual event. Boxplots indicate median (center line), interquartile ranges (25^th^-75^th^ percentile), and whiskers extending to 1.5×IQR. Inset (right) shows percent of events per duration subclass paired per-sample (dots) and group median (bars) ± IQR. **(H)** Event area; each dot is an individual event. Boxplots indicate median (center line), interquartile ranges (25^th^-75^th^ percentile), and whiskers extending to 1.5×IQR. Inset (right) shows percent of events per area subclass paired per-sample (dots) and group median (bars) ± IQR. **(I)** Forest plot summarizing treatment effects per parameter as percent change on a log_10_-scale. Amplitude, duration, and area show effect size and confidence intervals derived from linear mixed-effect models in black. Mean dF/F, frequency, and duration and area subclasses, show per-sample paired percent change across all samples as median with interquartile ranges (25^th^-75^th^ percentile) in blue (Wilcoxon signed-rank test). **(J)** Representative time-lapse images, **(K)** event dF/F traces, **(L)** mean fluorescence, **(M)** event amplitude, **(N)** raster plots, **(O)** event frequency, **(P)** event duration and sub-classes (inset), **(Q)** event area and sub-classes (inset), **(R)** and forest plot of human ioAstrocytes loaded with 2 µM Cal520-AM calcium dye and after treatment with 50 µM ATP. n = 23 (iAstrocytes) and 16 (ioAstrocytes) for respective ATP baseline-treatment pairs; * p > 0.05, ** p > 0.01, *** p > 0.001. Scale bar is 100 µm.

**Table 8:**
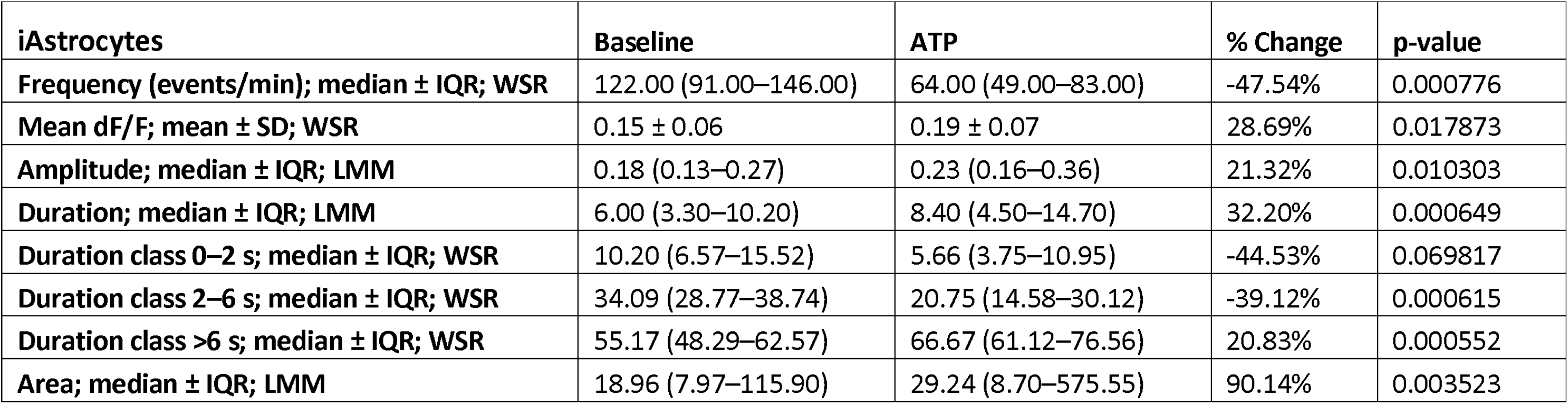
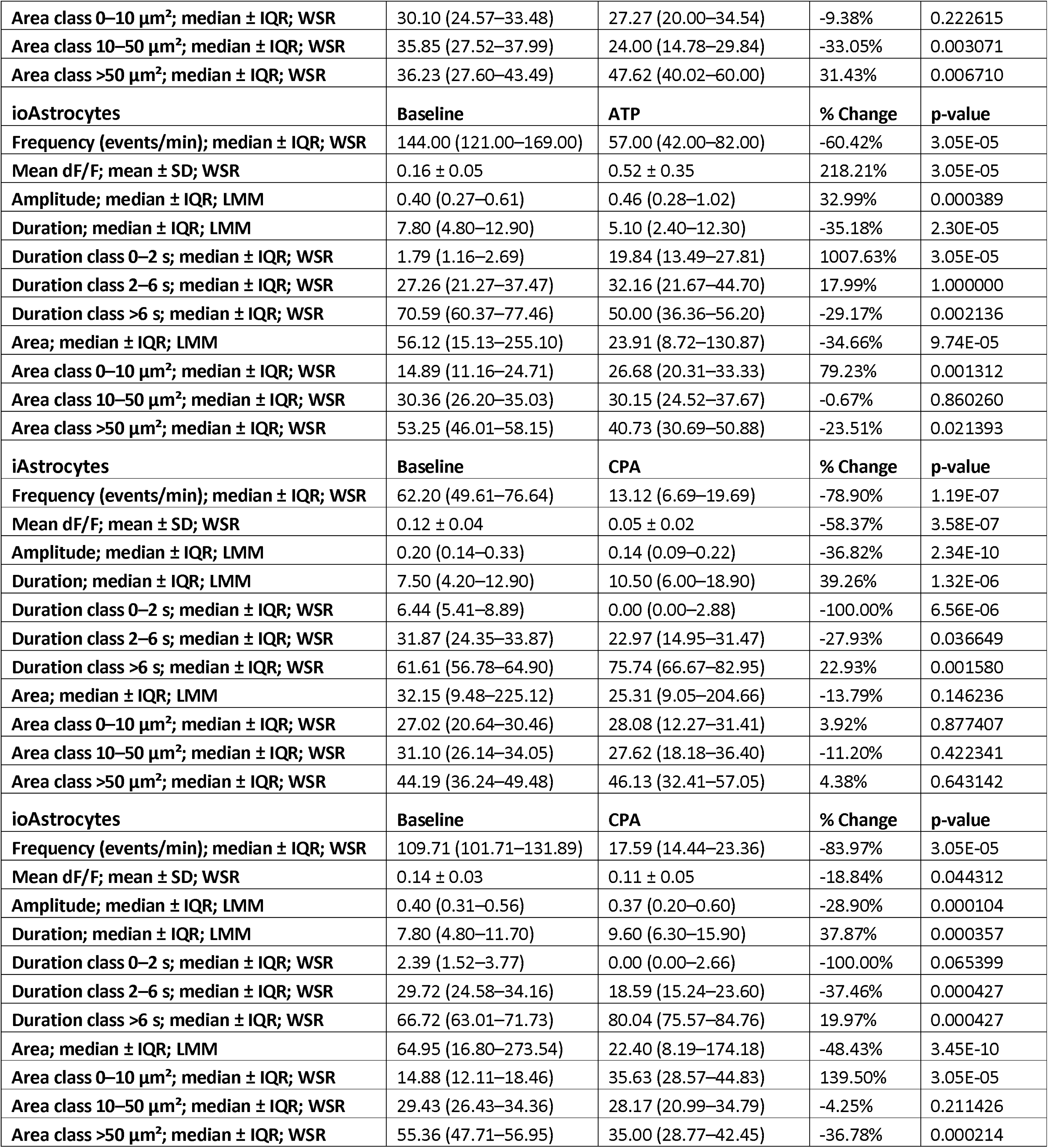
Descriptive statistics for effects of ATP and CPA in human iAstrocytes and human ioAstrocytes.

| **iAstrocytes** | **Baseline** | **ATP** | **% Change** | **p-value** |
| --- | --- | --- | --- | --- |
| **Frequency (events/min); median ± IQR; WSR** | 122.00 (91.00–146.00) | 64.00 (49.00–83.00) | -47.54% | 0.000776 |
| **Mean dF/F; mean ± SD; WSR** | 0.15 ± 0.06 | 0.19 ± 0.07 | 28.69% | 0.017873 |
| **Amplitude; median ± IQR; LMM** | 0.18 (0.13–0.27) | 0.23 (0.16–0.36) | 21.32% | 0.010303 |
| **Duration; median ± IQR; LMM** | 6.00 (3.30–10.20) | 8.40 (4.50–14.70) | 32.20% | 0.000649 |
| **Duration class 0–2 s; median ± IQR; WSR** | 10.20 (6.57–15.52) | 5.66 (3.75–10.95) | -44.53% | 0.069817 |
| **Duration class 2–6 s; median ± IQR; WSR** | 34.09 (28.77–38.74) | 20.75 (14.58–30.12) | -39.12% | 0.000615 |
| **Duration class >6 s; median ± IQR; WSR** | 55.17 (48.29–62.57) | 66.67 (61.12–76.56) | 20.83% | 0.000552 |
| **Area; median ± IQR; LMM** | 18.96 (7.97–115.90) | 29.24 (8.70–575.55) | 90.14% | 0.003523 |
| Area class 0–10 $\mu\text{m}^2$; median ± IQR; WSR | 30.10 (24.57–33.48) | 27.27 (20.00–34.54) | -9.38% | 0.222615 |
| Area class 10–50 $\mu\text{m}^2$; median ± IQR; WSR | 35.85 (27.52–37.99) | 24.00 (14.78–29.84) | -33.05% | 0.003071 |
| Area class >50 $\mu\text{m}^2$; median ± IQR; WSR | 36.23 (27.60–43.49) | 47.62 (40.02–60.00) | 31.43% | 0.006710 |
| **ioAstrocytes** | **Baseline** | **ATP** | **% Change** | **p-value** |
| Frequency (events/min); median ± IQR; WSR | 144.00 (121.00–169.00) | 57.00 (42.00–82.00) | -60.42% | 3.05E-05 |
| Mean dF/F; mean ± SD; WSR | 0.16 ± 0.05 | 0.52 ± 0.35 | 218.21% | 3.05E-05 |
| Amplitude; median ± IQR; LMM | 0.40 (0.27–0.61) | 0.46 (0.28–1.02) | 32.99% | 0.000389 |
| Duration; median ± IQR; LMM | 7.80 (4.80–12.90) | 5.10 (2.40–12.30) | -35.18% | 2.30E-05 |
| Duration class 0–2 s; median ± IQR; WSR | 1.79 (1.16–2.69) | 19.84 (13.49–27.81) | 1007.63% | 3.05E-05 |
| Duration class 2–6 s; median ± IQR; WSR | 27.26 (21.27–37.47) | 32.16 (21.67–44.70) | 17.99% | 1.000000 |
| Duration class >6 s; median ± IQR; WSR | 70.59 (60.37–77.46) | 50.00 (36.36–56.20) | -29.17% | 0.002136 |
| Area; median ± IQR; LMM | 56.12 (15.13–255.10) | 23.91 (8.72–130.87) | -34.66% | 9.74E-05 |
| Area class 0–10 $\mu\text{m}^2$; median ± IQR; WSR | 14.89 (11.16–24.71) | 26.68 (20.31–33.33) | 79.23% | 0.001312 |
| Area class 10–50 $\mu\text{m}^2$; median ± IQR; WSR | 30.36 (26.20–35.03) | 30.15 (24.52–37.67) | -0.67% | 0.860260 |
| Area class >50 $\mu\text{m}^2$; median ± IQR; WSR | 53.25 (46.01–58.15) | 40.73 (30.69–50.88) | -23.51% | 0.021393 |
| **iAstrocytes** | **Baseline** | **CPA** | **% Change** | **p-value** |
| Frequency (events/min); median ± IQR; WSR | 62.20 (49.61–76.64) | 13.12 (6.69–19.69) | -78.90% | 1.19E-07 |
| Mean dF/F; mean ± SD; WSR | 0.12 ± 0.04 | 0.05 ± 0.02 | -58.37% | 3.58E-07 |
| Amplitude; median ± IQR; LMM | 0.20 (0.14–0.33) | 0.14 (0.09–0.22) | -36.82% | 2.34E-10 |
| Duration; median ± IQR; LMM | 7.50 (4.20–12.90) | 10.50 (6.00–18.90) | 39.26% | 1.32E-06 |
| Duration class 0–2 s; median ± IQR; WSR | 6.44 (5.41–8.89) | 0.00 (0.00–2.88) | -100.00% | 6.56E-06 |
| Duration class 2–6 s; median ± IQR; WSR | 31.87 (24.35–33.87) | 22.97 (14.95–31.47) | -27.93% | 0.036649 |
| Duration class >6 s; median ± IQR; WSR | 61.61 (56.78–64.90) | 75.74 (66.67–82.95) | 22.93% | 0.001580 |
| Area; median ± IQR; LMM | 32.15 (9.48–225.12) | 25.31 (9.05–204.66) | -13.79% | 0.146236 |
| Area class 0–10 $\mu\text{m}^2$; median ± IQR; WSR | 27.02 (20.64–30.46) | 28.08 (12.27–31.41) | 3.92% | 0.877407 |
| Area class 10–50 $\mu\text{m}^2$; median ± IQR; WSR | 31.10 (26.14–34.05) | 27.62 (18.18–36.40) | -11.20% | 0.422341 |
| Area class >50 $\mu\text{m}^2$; median ± IQR; WSR | 44.19 (36.24–49.48) | 46.13 (32.41–57.05) | 4.38% | 0.643142 |
| **ioAstrocytes** | **Baseline** | **CPA** | **% Change** | **p-value** |
| Frequency (events/min); median ± IQR; WSR | 109.71 (101.71–131.89) | 17.59 (14.44–23.36) | -83.97% | 3.05E-05 |
| Mean dF/F; mean ± SD; WSR | 0.14 ± 0.03 | 0.11 ± 0.05 | -18.84% | 0.044312 |
| Amplitude; median ± IQR; LMM | 0.40 (0.31–0.56) | 0.37 (0.20–0.60) | -28.90% | 0.000104 |
| Duration; median ± IQR; LMM | 7.80 (4.80–11.70) | 9.60 (6.30–15.90) | 37.87% | 0.000357 |
| Duration class 0–2 s; median ± IQR; WSR | 2.39 (1.52–3.77) | 0.00 (0.00–2.66) | -100.00% | 0.065399 |
| Duration class 2–6 s; median ± IQR; WSR | 29.72 (24.58–34.16) | 18.59 (15.24–23.60) | -37.46% | 0.000427 |
| Duration class >6 s; median ± IQR; WSR | 66.72 (63.01–71.73) | 80.04 (75.57–84.76) | 19.97% | 0.000427 |
| Area; median ± IQR; LMM | 64.95 (16.80–273.54) | 22.40 (8.19–174.18) | -48.43% | 3.45E-10 |
| Area class 0–10 $\mu\text{m}^2$; median ± IQR; WSR | 14.88 (12.11–18.46) | 35.63 (28.57–44.83) | 139.50% | 3.05E-05 |
| Area class 10–50 $\mu\text{m}^2$; median ± IQR; WSR | 29.43 (26.43–34.36) | 28.17 (20.99–34.79) | -4.25% | 0.211426 |
| Area class >50 $\mu\text{m}^2$; median ± IQR; WSR | 55.36 (47.71–56.95) | 35.00 (28.77–42.45) | -36.78% | 0.000214 |

Application of ATP induced a rapid transient activation of ioAstrocytes (Fig. 8J), which was also observed in fluorescent traces (Fig. 8K, Movie 6). Mean fluorescence was increased within the first 30 seconds (Fig. 8L; *p*<0.001), consistent with all previous ATP experiments, where an increase in mean fluorescence was the most robust effect of ATP. Event amplitude was also increased (Fig. 8M; β=1.33, 95%-CI: 1.34, 1.56, *p*<0.001) as in iAstrocytes. Raster plots (Fig. 8N) indicate that, like iAstrocytes, ioAstrocytes tend to have longer events than mouse astrocytes. Furthermore, like iAstrocytes, a reduction in event frequency was observed in ioAstrocytes (Fig. 8O; *p*<0.001). Event duration decreased in ioAstrocytes after ATP application (Fig. 8P; β=0.65, 95%-CI: 0.53, 0.79, *p*<0.001) due to more 0-2 second events (*p*<0.001) and fewer >6 second events (*p*=0.0021). In contrast, event duration was increased in iAstrocytes and mouse astrocytes. Event area was decreased (Fig. 8Q; β=0.65, 95%-CI: 0.53, 0.81, *p*<0.001) due to a shift towards more microdomain activity (*p*=0.0013) and fewer wave events (*p*=0.021). This contrasted results from iAstrocytes, where not only the overall area, but also the proportion of waves increased. Statistics are summarized in Figure 8R and Table 8. In sum, ATP robustly increased mean fluorescence and event amplitude, and reduced event frequency in both human astrocyte cultures. Compounds targeting either of these parameters could therefore be tested in both iAstrocytes and ioAstrocytes. However, event duration and area showed opposing results.

### Human iPSC-derived astrocyte calcium signaling is decreased by CPA

Next, we tested if CPA would decrease activity in the two human astrocyte preparations as it did in mouse astrocytes. We observed a strong decrease in calcium signal in iAstrocytes (Fig. 9A) and in fluorescence traces (Fig. 9B) following application of CPA, confirmed by a decrease in mean fluorescence (Fig. 9C; *p*<0.001). This was in contrast to mouse astrocytes, where mean fluorescence increased. Event amplitude decreased after CPA application in iAstrocytes (Fig. 9D; β=0.63, 95%-CI: 0.55, 0.73, *p*<0.001), similar to mouse astrocytes in monoculture, but unlike cocultured mouse astrocytes, where amplitude increased. Raster plots (Fig. 9E) showed a clear reduction in event frequency (Fig. 9F; *p*<0.001), confirming the inhibitory effect of CPA also seen in mouse astrocytes. CPA increased event duration in iAstrocytes (Fig. 9G; β=1.40, 95%-CI: 1.22, 1.60, *p*<0.001) due to a shift from fast to slow events, with decreases in 0-2 (*p*<0.001) and 2-6 second events (*p*=0.037), and an increase in events >6 seconds (*p*=0.0016). This was similar to mouse astrocytes, with an increase in >6 second events being the most consistent effect of CPA across all datasets. A reduction in event area, as seen in mouse astrocytes, was not observed in iAstrocytes (Fig. 9H). Statistics are summarized in Figure 9I and Table 8.

**Figure 9:**
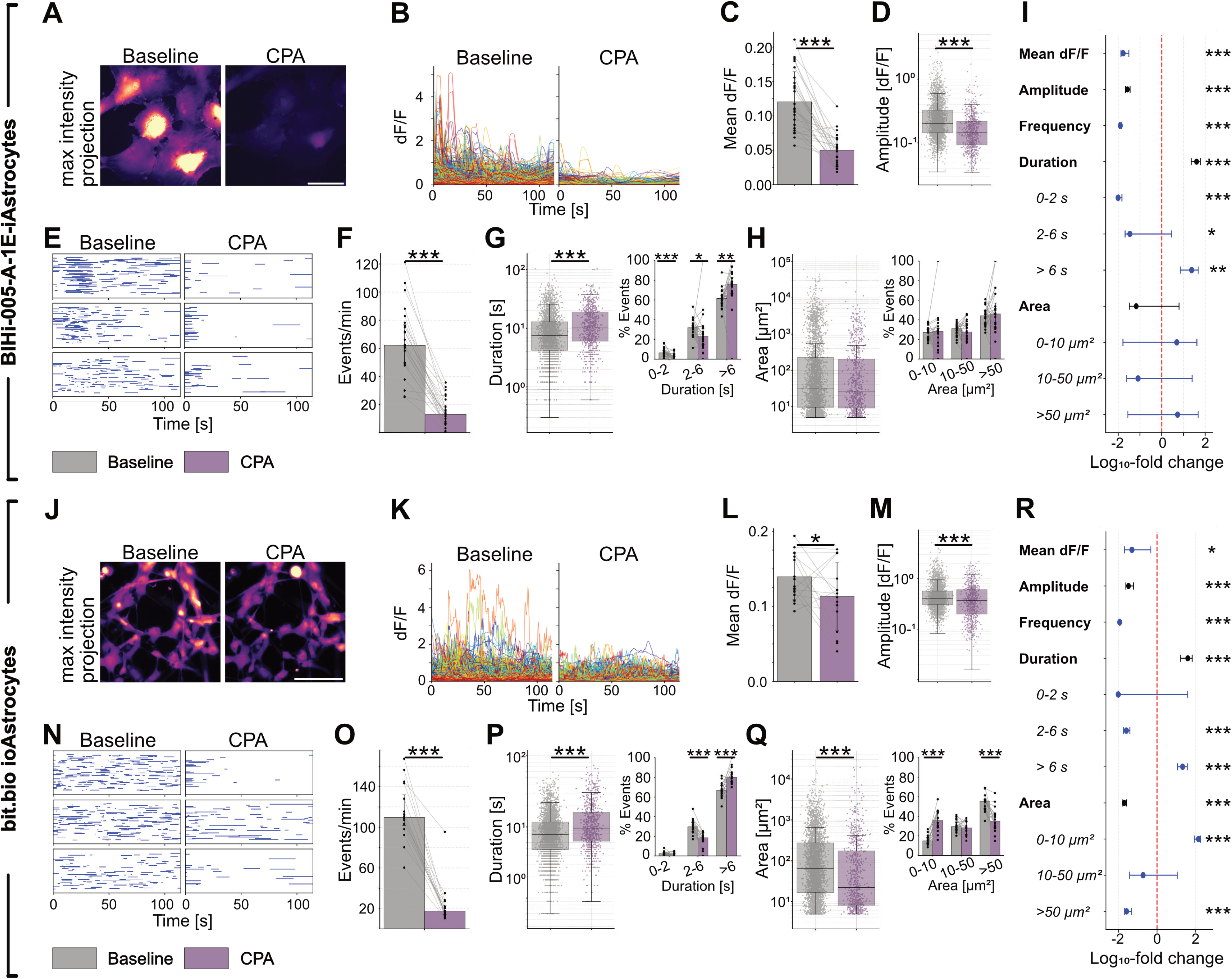
CPA decreased calcium activity in human iAstrocytes and ioAstrocytes. **(A)** Max intensity projections of a typical response induced by 20 µM CPA in human iAstrocytes loaded with 2 µM Cal520-AM calcium dye. **(B)** Individual event dF/F traces from baseline and treatment recordings. **(C)** Mean fluorescence per sample from event traces, displayed as paired data (dots) and group means (bars) ± SD. **(D)** Event amplitude; each dot is an individual event. Boxplots indicate median (center line), interquartile ranges (25^th^-75^th^ percentile), and whiskers extending to 1.5×IQR. **(E)** Raster plots showing occurrence of individual astrocytic events as blue horizontal lines, with length corresponding to event duration. **(F)** Event frequency, displayed as paired regions (dots) and group median (bars) ± IQR. **(G)** Event duration; each dot is an individual event. Boxplots indicate median (center line), interquartile ranges (25^th^-75^th^ percentile), and whiskers extending to 1.5×IQR. Inset (right) shows percent of events per duration subclass paired per-sample (dots) and group median (bars) ± IQR. **(H)** Event area; each dot is an individual event. Boxplots indicate median (center line), interquartile ranges (25^th^-75^th^ percentile), and whiskers extending to 1.5×IQR. Inset (right) shows percent of events per area subclass paired per-sample (dots) and group median (bars) ± IQR. **(I)** Forest plot summarizing treatment effects per parameter as percent change on a log_10_-scale. Amplitude, duration, and area show effect size and confidence intervals derived from linear mixed-effect models in black. Mean dF/F, frequency, and duration and area subclasses, show per-sample paired percent change across all samples as median with interquartile ranges (25^th^-75^th^ percentile) in blue (Wilcoxon signed-rank test). **(J)** Representative max intensity projections, **(K)** event dF/F traces, **(L)** mean fluorescence, **(M)** event amplitude, **(N)** raster plots, **(O)** event frequency, **(P)** event duration and sub-classes (inset), **(Q)** event area and sub-classes (inset), **(R)** and forest plot of human ioAstrocytes loaded with 2 µM Cal520-AM calcium dye and after treatment with 20 µM CPA. n = 24 (iAstrocytes) and 16 (ioAstrocytes) for respective CPA baseline-treatment pairs; * p > 0.05, ** p > 0.01, *** p > 0.001. Scale bar is 100 µm.

In ioAstrocytes, CPA decrease activity as seen in maximum intensity projections (Fig. 9J) and fluorescence traces (Fig. 9K), with a decrease in mean fluorescence (Fig. 9L; *p*=0.044) and event amplitude (Fig. 9M; β=0.71, 95%-CI: 0.60, 0.84, *p*<0.001) like iAstrocytes. Raster plots showed lower event densities (Fig. 9N), due to a decrease in event frequency (Fig. 9O; *p*<0.001), confirming the primary effect of CPA in ioAstrocytes. Event duration was increased (Fig. 9P; β=1.38, 95%-CI: 1.16, 1.64, *p*<0.001), with a decrease in 2-6 second events (*p*<0.001), and an increase in >6 second events (*p*<0.001), similar to iAstrocytes. Event area was decreased after application of CPA (Fig. 9Q; β=0.52, 95%-CI: 0.42, 0.63, *p*<0.001) with an increase in microdomain activity (*p*<0.001) and decrease in waves (*p*<0.001). This contrasts iAstrocytes, where CPA did not affect event area at all, but resembles effects seen in mouse astrocytes. Statistics are summarized in Figure 9R and Table 8. Between the two human astrocyte cultures, effects of CPA on mean fluorescence, event amplitude, frequency, and duration are robust and comparable, with only area being unchanged in iAstrocytes, and reduced in ioAstrocytes.

### LSD elicits opposing effects in human astrocytes than in cocultured mouse astrocytes

To evaluate the presence of functional serotonin receptors and the effects of LSD, we tested the effects of one-hour incubation of LSD on human ioAstrocytes. We observed increased fluorescence in response to LSD in maximum intensity projections (Fig. 10A), fluorescence traces (Fig. 10B), and mean fluorescence (Fig. 10C; *p*=0.048), in contrast to the decrease we observed in mouse astrocytes. Like in mouse astrocytes, event amplitude (Fig. 10D) remained unchanged. Raster plots showed higher event density (Fig. 10E) with increased event frequency after LSD application (Fig. 10F; *p*=0.0054), in contrast to the slight decrease observed in mouse cocultures. Event duration increased (Fig. 10G; β=1.19, 95%-CI: 1.09, 1.29, *p*<0.001), with fewer 2-6 second events (*p*=0.0012) and more >6 second events (*p*<0.001). No change in event area was observed (Fig. 10H), similar to cocultured mouse astrocytes. Statistics are summarized in Figure 10I and Table 9. In summary, LSD increased calcium signaling in human ioAstrocytes, opposite to LSD effects on cocultured mouse astrocytes. Importantly, the robust calcium response in ioAstrocytes demonstrates that they express functional serotonin receptors, supporting their use to test astrocytic responses to serotonergic compounds.

**Figure 10:**
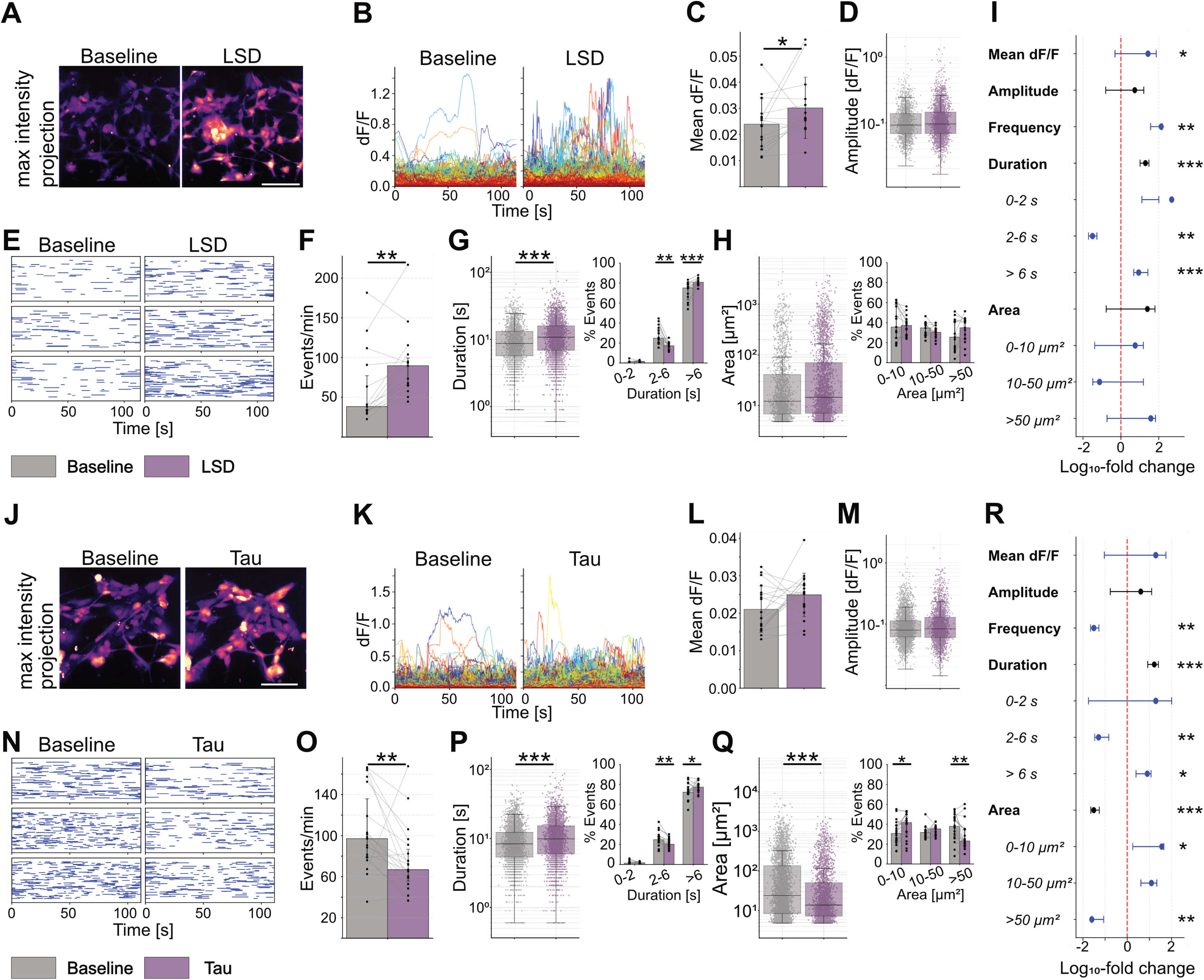
Effects of LSD and human Tau oligomers in human ioAstrocytes. **(A)** Max intensity projections of a typical response induced by 10 µM LSD after one hour in human ioAstrocytes loaded with 1 µM Cal520-AM calcium dye. **(B)** Individual event dF/F traces from baseline and treatment recordings. **(C)** Mean fluorescence per sample from event traces, displayed as paired data (dots) and group means (bars) ± SD. **(D)** Event amplitude; each dot is an individual event. Boxplots indicate median (center line), interquartile ranges (25^th^-75^th^ percentile), and whiskers extending to 1.5×IQR. **(E)** Raster plots showing occurrence of individual astrocytic events as blue horizontal lines, with length corresponding to event duration. **(F)** Event frequency, displayed as paired regions (dots) and group median (bars) ± IQR. **(G)** Event duration; each dot is an individual event. Boxplots indicate median (center line), interquartile ranges (25^th^-75^th^ percentile), and whiskers extending to 1.5×IQR. Inset (right) shows percent of events per duration subclass paired per-sample (dots) and group median (bars) ± IQR. **(H)** Event area; each dot is an individual event. Boxplots indicate median (center line), interquartile ranges (25^th^-75^th^ percentile), and whiskers extending to 1.5×IQR. Inset (right) shows percent of events per area subclass paired per-sample (dots) and group median (bars) ± IQR. **(I)** Forest plot summarizing treatment effects per parameter as percent change on a log_10_-scale. Amplitude, duration, and area show effect size and confidence intervals derived from linear mixed-effect models in black. Mean dF/F, frequency, and duration and area subclasses, show per-sample paired percent change across all samples as median with interquartile ranges (25^th^-75^th^ percentile) in blue (Wilcoxon signed-rank test). **(J)** Representative max intensity projections, **(K)** event dF/F traces, **(L)** mean fluorescence, **(M)** event amplitude, **(N)** raster plots, **(O)** event frequency, **(P)** event duration and sub-classes (inset), **(Q)** event area and sub-classes (inset), **(R)** and forest plot of human ioAstrocytes loaded with 1 µM Cal520-AM calcium dye after one hour treatment with 25 nM human Tau oligomers. n = 15 (LSD) and 19 (Tau) for respective baseline-treatment pairs; * p > 0.05, ** p > 0.01, *** p > 0.001. Scale bar is 100 µm.

**Table 9:** Descriptive statistics for effects of LSD and human Tau in human ioAstrocytes.

|  | Baseline | LSD | % Change | p-value |
| --- | --- | --- | --- | --- |
| Frequency (events/min); median ± IQR; WSR | 38.32 (33.86–77.17) | 89.76 (65.09–99.48) | 134.25% | 0.005371 |
| Mean dF/F; mean ± SD; WSR | 0.02 ± 0.01 | 0.03 ± 0.01 | 26.06% | 0.047913 |
| Amplitude; median ± IQR; LMM | 0.09 (0.07–0.14) | 0.10 (0.07–0.15) | 4.42% | 0.387331 |
| Duration; median ± IQR; LMM | 8.70 (5.70–13.20) | 10.80 (6.90–15.90) | 18.93% | 5.34E-05 |
| Duration class 0–2 s; median ± IQR; WSR | 0.29 (0.00–1.41) | 1.69 (0.85–2.18) | 483.15% | 0.055359 |
| Duration class 2–6 s; median ± IQR; WSR | 24.88 (20.39–33.78) | 17.27 (14.92–19.94) | -30.58% | 0.001225 |
| Duration class >6 s; median ± IQR; WSR | 75.12 (63.21–78.84) | 80.91 (78.63–83.29) | 7.71% | 0.000854 |
| Area; median ± IQR; LMM | 12.18 (6.83–40.97) | 14.60 (7.04–69.96) | 24.18% | 0.111860 |
| Area class 0–10 $\mu\text{m}^2$; median ± IQR; WSR | 35.59 (24.70–57.66) | 37.27 (27.36–42.19) | 4.72% | 0.638672 |
| Area class 10–50 $\mu\text{m}^2$; median ± IQR; WSR | 34.71 (27.54–39.14) | 30.41 (27.08–33.56) | -12.39% | 0.276855 |
| Area class >50 $\mu\text{m}^2$; median ± IQR; WSR | 25.42 (14.46–41.02) | 34.88 (23.91–41.82) | 37.21% | 0.063721 |
|  | **Baseline** | **human Tau** | **% Change** | **p-value** |
| Frequency (events/min); median ± IQR; WSR | 97.04 (78.98–135.85) | 66.85 (55.80–86.79) | -31.11% | 0.002399 |
| Mean dF/F; mean ± SD; WSR | 0.02 ± 0.01 | 0.02 ± 0.01 | 18.39% | 0.104156 |
| Amplitude; median ± IQR; LMM | 0.08 (0.06–0.11) | 0.08 (0.06–0.13) | 3.07% | 0.458056 |
| Duration; median ± IQR; LMM | 8.40 (5.40–12.30) | 9.90 (6.00–15.30) | 15.63% | 0.000117 |
| Duration class 0–2 s; median ± IQR; WSR | 1.36 (0.44–2.25) | 1.61 (0.78–2.70) | 18.44% | 0.647471 |
| Duration class 2–6 s; median ± IQR; WSR | 24.80 (21.92–28.10) | 20.18 (17.75–24.07) | -18.61% | 0.003918 |
| Duration class >6 s; median ± IQR; WSR | 72.19 (70.89–76.09) | 77.27 (74.24–80.05) | 7.05% | 0.014069 |
| Area; median ± IQR; LMM | 23.95 (8.40–135.37) | 13.76 (7.25–50.47) | -31.26% | 7.67E-05 |
| Area class 0–10 $\mu\text{m}^2$; median ± IQR; WSR | 30.77 (23.24–38.78) | 41.62 (30.04–47.62) | 35.28% | 0.020412 |
| Area class 10–50 $\mu\text{m}^2$; median ± IQR; WSR | 31.69 (27.31–33.48) | 35.29 (29.52–37.07) | 11.36% | 0.054573 |
| Area class >50 $\mu\text{m}^2$; median ± IQR; WSR | 37.88 (28.43–46.56) | 23.21 (18.46–34.67) | -38.71% | 0.002838 |

### Tau-dependent calcium signaling changes in human astrocytes

Next, we tested the response of human ioAstrocytes to Tau oligomers. Tau oligomer application for one hour slightly increased fluorescence in maximum projection images (Fig. 10J) and in fluorescence traces (Fig. 10K), with no change in mean fluorescence (Fig. 10L) or event amplitude (Fig. 10M). Raster plots (Fig. 10N) indicated a decrease in event frequency (Fig. 10O; *p*=0.0024). All of these effects were similar to those observed in mouse astrocytes. Event duration increased after Tau application (Fig. 10P; β=1.16, 95%-CI: 1.07, 1.24, *p*<0.001) with fewer 2-6 second events (*p*=0.0039) and more >6 second events (*p*=0.014). In contrast, cocultured mouse astrocytes displayed a decrease in event duration. Event area was decreased (Fig. 10Q; β=0.69, 95%-CI: 0.57, 0.83, *p*<0.001), due to an increase in microdomain activity (*p*=0.020) and a decrease in waves (*p*=0.0028), similar to cocultured mouse astrocytes. Statistics are summarized in Figure 10R and Table 9. In summary, the effects of Tau oligomers on calcium event frequency, amplitude, and area, but not event duration, were robust across mouse and human astrocytes.

## DISCUSSION

We demonstrate that our calcium imaging and analysis pipeline can be used to characterize compound-specific effects on multiple parameters of calcium signaling in different astrocyte culture systems in formats up to 384-well plates. We benchmarked stimulatory and inhibitory effects of ATP and CPA, probed glutamatergic, serotonergic, and GABAergic receptor function, found novel effects of LSD in human astrocytes, and tested a Tau disease model. Calcium signals were sensitive to species (mouse vs. human), the presence or absence of neurons, and different human iPSC lines used for astrocyte induction. Our pipeline quantifies multiple fundamental parameters following event-based detection, ensuring that any effect on astrocytic calcium signaling is captured. This is useful for quickly and easily determining astrocytic effects of astrocyte-targeting compounds, off-target/secondary effects of compounds, or astrocytic effects of disease models – despite limited knowledge regarding the interpretation of astrocyte calcium data.

It is often unclear whether increases or decreases in specific astrocytic calcium signaling parameters are beneficial or detrimental, and what underlies these changes. However, the duration of calcium event fluorescence traces may give insights into the nature of activated receptors. Event duration increases if signals last longer, which could correspond to GPCR-dependent receptor types, while shorter events could indicate rapid calcium influx through ionotropic receptors. P2X and P2Y receptors are upregulated in the presence of neurons (Hasel et al., 2017), which may lead to differences in ATP responses in astrocytes in monocultures compared to cocultures, for example. Interestingly, mild ischemic injury in mice activates astrocytes and increases expression of P2X7 receptors, which are less responsive to ATP and thus reduce brain damage after middle cerebral artery occlusion (Hirayama et al., 2015).

Changes in certain astrocytic calcium signalling parameters could reflect pathological states. For example, a mouse model of Alexander’s disease, a rare genetic disorder caused by a mutation in the GFAP gene (Messing et al., 2012), exhibits dramatically increased area of calcium transients (Saito et al., 2018). Thus, a compound that reliably decreases event area might be a potential candidate for the treatment of Alexander’s disease. Astrocytes carrying the mutation could be tested for compounds that normalize event area. It is thought that the changes in event area in Alexander’s disease are caused by a dysfunction of the SERCA pump on the endoplasmic reticulum (Saito et al., 2018). Thus, compounds that restore calcium pump function could therefore be selected.

Both monocultures and cocultures of astrocytes are important models to study the properties of a compound. Monocultures provide information about astrocyte-specific effects of a compound: if the compound triggers a change of activity, it affects astrocytes. On the other hand, testing a drug that was specifically designed to target neurons in astrocyte monocultures would identify possible off-target effects. This comes with the caveat that astrocytes in monoculture differ in morphology and gene expression from cocultured astrocytes. Therefore, it is also important that compounds are evaluated in coculture systems where astrocyte activity is recorded. However, even a neuron-specific compound could lead to changes in calcium activity in astrocytes and indicate a secondary effect through the neurons. Thereby, astrocyte function could be an additional readout for neuron-targeted compounds. This could even reveal subtle compound effects that do not change neuronal activity but induce, for example, altered release of signaling molecules from neurons to astrocytes.

Using our pipeline, we made the novel observation that LSD affects astrocytic calcium signaling – but LSD exerts opposite effects on mouse and human astrocytes. In mouse cocultures, LSD decreased mean fluorescence and event frequency. However, application of LSD to human ioAstrocytes increased mean fluorescence, event frequency, and duration. Both results demonstrate an effect of LSD on astrocyte calcium dynamics, though in different directions. This could be due to either the presence of neurons and thus a neuronal effect on the astrocytes or to species differences. To date, research on LSD has focused on neuronal effects and synaptic plasticity (Costa et al., 2024; Ly et al., 2018; Ornelas et al., 2022), where LSD enhances neuronal plasticity via 5HT serotonergic receptors. LSD resulted in an antidepressant effect in rats, while this effect was absent in mice (Bouloufa et al., 2025), highlighting that even closely related species respond differently. LSD is indeed a potential therapeutic agent for major depressive disorders with successful outcomes in small clinical trials (Ko et al., 2023; Müller et al., 2025). Astrocyte calcium activity in a corticosterone-induced model of depression in mice is characterised by a decreased event duration (González-Arias et al., 2023). Intriguingly, LSD reliably increased event duration in human, but not in mouse astrocytes in our hands, suggesting LSD treatment could reverse alterations in calcium seen in depression models, if human models of depression also have decreased event duration. Both the reduced calcium signaling in mouse astrocytes and the increased signaling in human astrocytes after LSD treatment are, to our knowledge, novel findings. Further experiments are needed to characterize the effects of LSD under different treatment durations to reveal short- or long-term changes in astrocyte calcium dynamics induced by LSD. However, our experiments demonstrate the presence of functional serotonin receptors on astrocytes. Thus, astrocytes are a target for serotonin receptor targeting compounds. Astrocytes express functional 5-HT2A, the primary receptor for LSD, on their membrane (Hagberg et al., 1998), along with other serotonin receptor subtypes such as 5-HT1A (Whitaker-Azmitia et al., 1993), 5-HT4R (Müller et al., 2021), 5-HT7 (Shimizu et al., 1996), and serotonin transporters (Inazu et al., 2001). Both 5-HT and the selective-serotonin reuptake inhibitors fluoxetine and citalopram trigger calcium responses in astrocytes (Jalonen et al., 1997; Schipke et al., 2011). Activation of astrocytic, but not neuronal, 5-HT_4_R increased miniature excitatory postsynaptic potentials in neurons in vitro, and decreased paired-pulse ratio in situ, indicating modulatory effects of astrocytes on neurons (Müller et al., 2021). This could have positive effects on other diseases and disorders, in addition to depression. For example, LSD might be suitable to treat Alzheimer’s as it normalizes cortisol and glucocorticoid receptor-induced hyperstimulation of astrocytes (Shlomi et al., 2020).

Tau oligomer-treated astrocytes showed similar responses in both mouse and human astrocytes, with the primary effects observed being reduced event frequency and area. This observation is similar to previous data, in which exogenous oligomeric recombinant human Tau reduced frequency and amplitude of ATP-induced calcium events in astrocyte monocultures (Piacentini et al., 2017). Reduced astrocyte calcium event frequency was also found in an APP overexpression Alzheimer’s mouse model (Shah et al., 2022). Thus, a reduction in calcium event frequency could be an astrocytic calcium signaling signature associated with Alzheimer’s disease. The Tau oligomer model could be useful for testing calcium dynamics in astrocytes in response to other Tau species or durations of application. More importantly, the presented pipeline could be used to screen for compounds that rescue Tau oligomer-induced deficits in astrocyte calcium signaling, and hence help to discover new drug candidates that can prevent or reverse the pathological effects of Tau on astrocytes.

Our pipeline could be used in other disease models, to test effects on astrocytic calcium signaling and their rescue. For example, PS2APP Alzheimer model mice have reduced astrocytic calcium signaling due to reduced expression of the ER protein STIM1 that mediates store-operated calcium entry. Lentiviral expression of STIM1 normalized astrocytic calcium activity as well as performance in memory tasks (Lia et al., 2023). Moreover, AAP/PS1 mice exhibit impaired slow wave oscillations and astrocyte calcium transients that could be normalized by optogenetic activation of astrocytes, which also restored neuron function and memory (Lee et al., 2023). This illustrates that aberrant astrocytic calcium signaling is a disease signature that when restored has the potential to rescue disease phenotypes. Our pipeline could be used to characterize specific perturbed parameters in disease models and then screen for compounds that rescue them.

Astrocyte calcium signals are still insufficiently studied. Disease-related models may provide indications of “good” or “bad” patterns of activity. However, the direction of changes (increase or decrease in astrocytic calcium signaling) varies between diseases. In a study of autism spectrum disorder, for example, calcium signals were changed in opposite directions compared to baseline, and could therefore only be described as “aberrant” (Allen et al., 2022). Our pipeline is therefore designed to provide information on effects of all aspects of astrocyte calcium signaling perturbation in any direction. Using the pipeline, especially with human astrocytes, could improve drug testing by incorporating astrocytic effects of compounds, including new therapeutic approaches directly targeting astrocyte function.

## DATA AVAILABILITY

Data analyzed in this paper and the Python code used for analysis are available from the corresponding author upon request.

## AUTHOR CONTRIBUTIONS

JK and CD designed the study. LB prepared and supported the use of the BIHi-005-A-1E hiPSC-derived human astrocytes. SW prepared and provided human recombinant Tau oligomers. JK collected and analyzed the data. JK and CD wrote and edited the manuscript. CD provided funding and materials.

## ACKNOWLEDGEMENTS

We thank bit.bio for continuous support with the ioAstrocyte cultures, the BIH Core Unit pluripotent Stem Cells and Organoids (CUSCO) for the provision of BIHi005-A-1E derived human iAstrocytes, and the Charité Advanced Medical Bioimaging (AMBIO) core facility for microscopy support.

